# Identification and inhibition of the Cyclin D Rb-docking interface that drives cell division

**DOI:** 10.64898/2026.01.14.699544

**Authors:** Benjamin R. Topacio, Cecelia Brown Fleming, Michael C. Lanz, Mihkel Örd, Mina Memarpoor-Yazdi, Shuyuan Zhang, Shicong Xie, Jurgen Tuvikene, Aiden Weaver, Ioannis Sanidas, Norman E Davey, Peter M Pryciak, Julien Sage, Seth M. Rubin, Mardo Kõivomägi, Jan M. Skotheim

## Abstract

The animal cell division cycle is initiated by the cyclin-dependent kinases CDK4 and CDK6 in complex with D-type cyclins. Cyclin D-CDK4/6 complex formation is promoted by the assembly factors p21 and p27, which bind both subunits. p27 binds the hydrophobic patch on cyclin D that is similar to the patch used by other cell cycle cyclins to dock their substrates. This raised the question as to how cyclin D could find its substrates if its hydrophobic patch were already occupied? Here, we show that D-type cyclins use their A2’ helix to dock the retinoblastoma protein Rb, a key substrate regulating cell cycle progression. The specific interface of cyclin D’s A2’ helix is unique among cyclins, and mutation or inhibition of the A2’ helix slows proliferation. Taken together, our work identifies a cyclin D-substrate docking mechanism that can be targeted by novel cancer therapeutics.

## INTRODUCTION

The decision to transition from G1 to S phase of the cell cycle is critical to normal development and its misregulation is a hallmark of cancer. During G1, cells decide to either remain in G1 and wait for additional signals, to exit the cell cycle and enter quiescence, senescence or differentiation, or to proceed through the G1/S transition, replicate their DNA, and divide. The multiple signals determining whether or not cells proliferate are integrated by the G1/S cell cycle pathway^1–3^.

The first step in the G1/S transition is driven by growth factors triggering the synthesis of D-type cyclins (cyclin D1, D2, and D3), which form complexes with the cyclin-dependent kinases CDK4 and CDK6 to create the upstream activators of the G1/S transition^4,5^. The activity of cyclin D-CDK4/6 kinases is modulated by the levels of Cyclin D and also by CDK inhibitor proteins of the INK4 and CIP/KIP families, whose levels are controlled by cellular stress and oncogenic signals^6^. Somewhat paradoxically, cyclin D-CDK4/6 activity is also promoted by the CIP/KIP proteins p21 and p27, which are generally considered cell cycle inhibitors. However, p21 and p27 also serve as assembly and activation factors for cyclin D-CDK4/6 complexes^6,7^.

Active cyclin D-CDK4/6 complexes phosphorylate and inhibit the function of the Rb family of cell cycle inhibitors. When unphosphorylated, Rb interacts with members of the E2F family of transcription factors to repress the transcription of genes whose products are critical for cell growth, DNA replication, and cell cycle progression^8–10^. Phosphorylation takes place on up to 15 sites and drives structural changes in Rb that lead to E2F dissociation^11^. While there is a consensus that cyclin D-CDK4/6 complexes phosphorylate Rb, the dynamics of this process have recently been contested^2,3^. The previous consensus was that cell growth in G1 drove a gradual increase in cyclin D-CDK4/6 activity which eventually activated the E2F-dependent transcription of the downstream cyclins E and A that complete Rb hyperphosphorylation^12–14^. This view was challenged by two newer, and competing, models. In the first new model, Rb phosphorylation is a two-step process, where upstream cyclin D-CDK4/6 mono-phosphorylates Rb to modulate its function before downstream cyclin E/A-CDK2 complexes complete Rb hyperphosphorylation ^15,16^. The ability of mono-phosphorylated Rb to repress E2F-dependent expression suggested that cyclin D phosphorylation of Rb did not promote the G1/S transition. However, our previous work shows that mutation of Rb to prevent cyclin D from docking resulted in a delayed G1/S transition and reduced Rb phosphorylation^17^. While this model does not have an obvious way to couple E2F-dependent expression of cyclins E and A to cell growth, this could be provided by growth diluting Rb in G1^18,19^. In a second model, cyclin D-CDK4/6 completes Rb hyperphosphorylation on its own, which is then maintained by the E2F-dependent synthesis of cyclins E and A^20,21^. In any case, while the dynamics and mechanisms of Rb phosphorylation are still contested, there is a consensus that Rb is a major target of cyclin D-CDK4/6.

While it is clear that Rb is phosphorylated by cyclin D-CDK4/6 complexes, our understanding of how these cyclin D-CDK4/6 complexes recognize Rb and other substrates has been incomplete. The cyclin D subunit is responsible for substrate docking and, to date, three distinct docking mechanisms have been reported: two support phosphorylation of Rb by CDK4/6, while the third is independent of CDK4/6 and does not engage kinase activity^17,22^. Of the two phosphorylation-linked mechanisms the first identified was an LxCxE motif on cyclin D that engages a corresponding binding cleft on Rb^23^. However, mutation of this motif produces only a modest reduction in cyclin D-CDK4/6 activity towards Rb *in vitro*, indicating that it is not the dominant docking mechanism^24^. Separately, the kinase-independent mechanism relies on an LLxxxL motif in the cyclin D C-terminus, which mediates interactions with transcription factors such as the estrogen receptor and SRC-1 to drive estrogen-responsive gene expression in a CDK- and estrogen-independent manner^25–27^. The second and more recently identified phosphorylation-linked mechanism, by contrast, involves recognition of the C-terminal helix of Rb through an unknown location on cyclin D. This helix-docking interaction appears to be the major substrate docking mechanism with respect to cell cycle control, as mutation of the relevant interface residues on Rb is sufficient to promote G1 arrest^24^. Yet, despite its clear importance for cell cycle progression, the molecular basis of this interaction on the cyclin D side remains unresolved (**Fig. 1A**).

**Figure 1:**
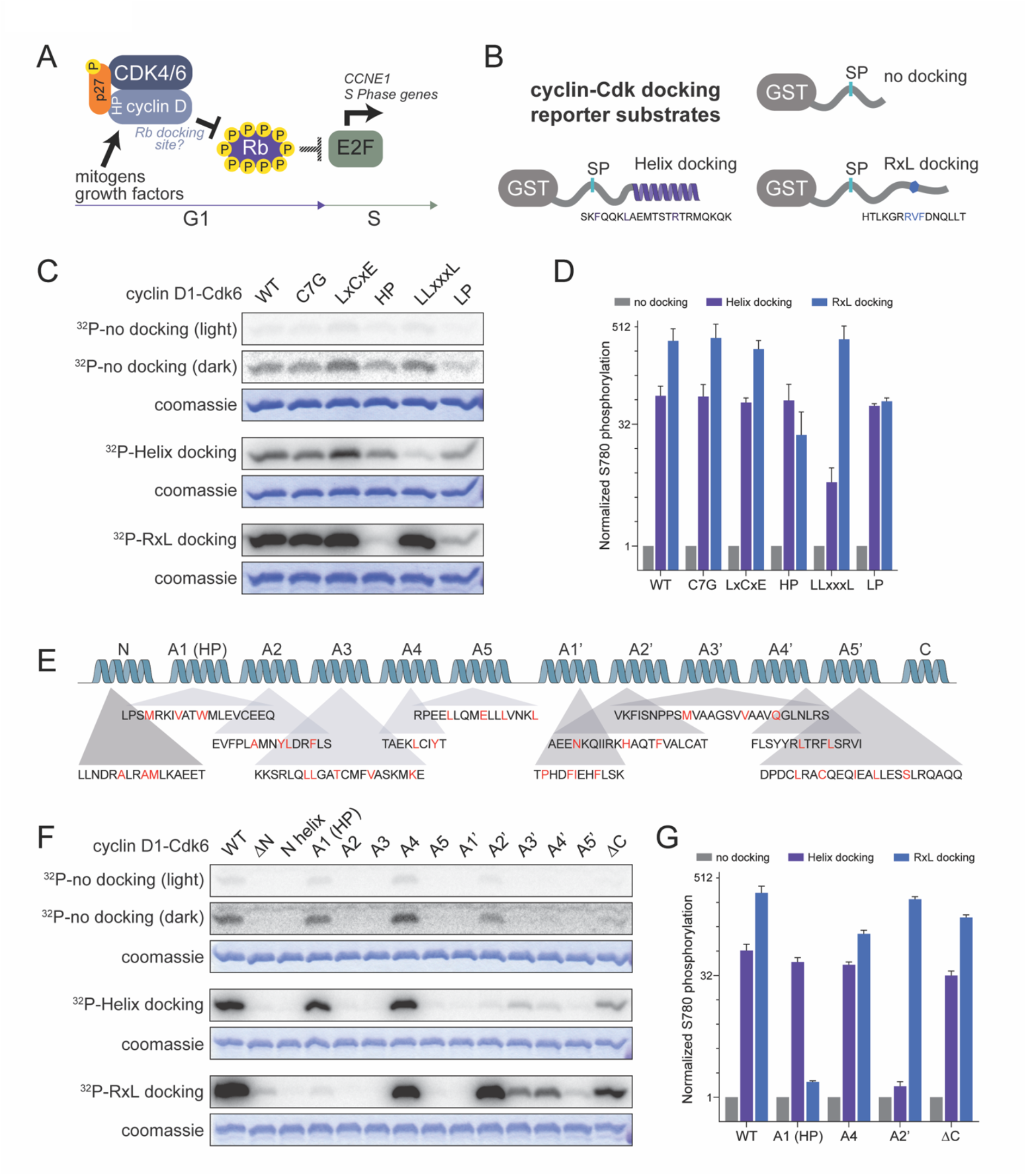
Cyclin D’s A2’ helix docks synthetic substrates containing Rb’s C-terminal helix. A. Schematic of the G1/S transition driven by cyclin D-CDK4/6. The kinase complex initiates E2F de-repression and S phase by phosphorylating Rb. Since the hydrophobic patch (HP) of active complexes is occupied by the p27 assembly factor, it is unclear what part of cyclin docks Rb. B. Schematic of the synthetic cyclin-Cdk substrate reporter fusion proteins containing a GST tag and the Rb amino acids 775-787 that include a single Cdk target phosphorylation site. These synthetic substrates are fused to a short amino acid sequence containing either no docking site, an Rb C-terminal helix docking site (Helix), or an RxL docking site derived from Cdc6 (RxL). C. Representative autoradiographs showing results from *in vitro* kinase assays using the indicated synthetic substrate and the indicated cyclin D1-Cdk6 kinase variant. WT denotes wild-type cyclin D1, C7G and LxCxE denote two different mutations of the LxCxE motif, HP denotes mutation of the hydrophobic patch, and LLxxxL denotes mutation of the steroid receptor-binding motif. The coomassie stained gels showing equal amounts of substrate used in each reaction are shown below each autoradiograph. D. Quantification of kinase assays from B (n=3). E. Schematic detailing the alpha helices on cyclin D1. The N-terminal cyclin box fold contains five alpha helices, A1-A5, and the C-terminal cyclin box fold contains the additional five alpha helices, A1’-A5’. The A1 helix contains the previously known hydrophobic patch groove (HP). The amino acids sequence for each helix is shown below the protein schematic and the interface residues potentially involved in Rb-docking that were substituted with alanines are colored red. F. Representative autoradiograph of *in vitro* kinase assays of the indicated synthetic substrate by the indicated kinase. WT denotes wild-type cyclin D1. The coomassie stained gels showing equal amounts of substrate used in each reaction are shown below each autoradiograph. We note that many of these cyclin D1-Cdk6 variants lack intrinsic kinase activity, as demonstrated by lack of signal for any substrate in the autoradiograph. G. Quantification of E (n=3).

All three D-type cyclins dock Rb’s C-terminal helix to drive Rb phosphorylation and cell cycle progression^24^, but it is unclear what part of cyclin D docks the Rb helix. Animal cyclin docking has been best studied for cyclins E and A, which have a hydrophobic patch that recognizes RxL motifs in unstructured parts of target proteins including p21 and p27^28,29^. Cyclin D also has a hydrophobic patch and we previously found a cyclin D-CDK4/6 dimer can dock RxL sequences, including those in Rb^24^. However, it is unlikely that cyclin D’s hydrophobic patch is available for docking substrates in cells because it binds an RxL motif in p27, which serves as an assembly factor for the cyclin D-CDK4,6 complex ^7^. Indeed, because CDK4 activity was low in cells lacking p27 and p21, it is likely that the CDK4 complex in cells is a trimer that also contains one of these assembly factors^30–33^. This suggests that cyclin D is capable of RxL-based docking through its hydrophobic patch but is unlikely to use this patch to dock its substrates in cells because it is already utilized to assemble active complexes with CDK4/6.

Here, we report the discovery and characterization of the interface on cyclin D that mediates helix-based docking with Rb to drive cell cycle progression. We screened cyclin D1 variants using *in vitro* kinase assays phosphorylating engineered CDK substrate proteins and identified the cyclin D interface that docks Rb’s C-terminal helix. This interface maps to cyclin D’s A2’ helix, and its mutation both reduces Rb phosphorylation and inhibits cell cycle progression. To directly test this interface, we targeted it with an Rb-based competitor peptide. Expression in cells of the wild-type Rb C-terminal peptide specifically inhibited proliferation and was synergistic with CDK4/6 inhibitor treatment. Taken together, our work has identified the two specific alpha helices on Rb and cyclin D whose interface defines the cyclin D-Rb docking mechanism driving the G1/S transition.

## Results

### Previously known sites on cyclin D do not dock Rb’s C-terminal helix

To determine the site on cyclin D that binds Rb’s C-terminal helix, we initially sought to test previously reported docking sites using *in vitro* kinase assays (see Methods). We anticipated that a cyclin D mutant lacking helix-based docking in complex with either CDK4 or CDK6 would satisfy three criteria. First, such a cyclin D mutant would have a baseline level of kinase activity against a non-docking substrate similar to wild-type. Second, the mutant would have an increased ability to phosphorylate an RxL-docking substrate over the non-docking substrate through its intact hydrophobic patch. And third, the mutant would lose the ability to phosphorylate helix-docking substrates (**Fig. 1B**). To test previously identified cyclin D docking sites, we generated cyclin D1 variants with alanine substitutions in the LxCxE motif (cyclin D1^C7G^ or cyclin D1^LxCxE^), the hydrophobic patch (cyclin D1^HP^), or the LLxxxL motif (cyclin D1^LLxxxL^)^23,26,27,34,35^. These cyclin D variants were fused to CDK6 to facilitate purification of active complexes as described previously^24^.

After generating a series of cyclin D1 variants fused to CDK6, we measured the activity of these complexes towards synthetic CDK substrate reporter proteins containing a GST tag and the Rb amino acids 775–790 containing a single CDK site (**Fig. 1C**). These synthetic substrates also contained an Rb C-terminal helix docking sequence, an RxL docking sequence, or no docking sequence. These variant enzymes maintained intrinsic kinase activity because they could phosphorylate the non-docking substrate (**Fig. 1C-D**). As we found previously, WT cyclin D1 complexes had much higher activity towards the helix-docking substrate than the non-docking substrate (71.5±13.2-fold increase). Similarly, cyclin D1^LxCxE^-CDK6 and cyclin D1^HP^-CDK6 complexes exhibited 59.1±4.8-fold and 63.0±21.1-fold higher activity towards the helix-docking substrate than the non-docking substrate, which was a similar increase as for the WT cyclin D1. Consistent with the expectation that the hydrophobic patch docks RxL sequences, mutation of the hydrophobic patch on the cyclin D1^HP^ variant resulted in lost activity towards the RxL-docking substrate. While cyclin D1^WT^-CDK6 showed 342.1±81.0-fold increased activity towards the RxL-docking substrate compared to the non-docking substrate, cyclin D1^HP^-CDK6 showed only a 23.6±12.5-fold increased activity. Cyclin D1^LLxxxL^-CDK6 showed 357.0±95.4-fold increased activity towards the RxL-docking substrate compared to the non-docking substrate, which was similar to cyclin D1^WT^-CDK6. Interestingly, cyclin D1^LLxxxL^-CDK6 activity towards the helix-docking substrate was partially impaired and showed only a 6.1±1.8-fold increase relative to activity towards the non-docking substrate. This suggests that the LLxxxL motif may contribute the helix-based docking on cyclin D1 but is unlikely to be solely responsible, particularly given that this motif is not conserved among the D-type cyclins^17^. Another possibility is that the LLxxxL mutation resulted in a slight structural change affecting the position of the actual docking interface to reduce its effectiveness. Since we expect that mutation of the helix-docking interface would result in a larger loss in activity towards the helix-docking reporter substrate, we continued our search for a cyclin D1 mutant that disrupts helix-based docking.

Having examined the previously known docking sites on cyclin D1, we next sought to examine docking sites identified on other cyclins because of the known structural homology of the cyclin family of proteins. In budding yeast, a helix in the G1 cyclin Cln2 docks a linear LP motif in its substrates, including Whi5, the functional ortholog of Rb^36^. To test if the homologous site on cyclin D1 was responsible for helix-based docking, we generated a cyclin D1^LP^-CDK6 complex that has a mutation in this potential G1 cyclin docking site. However, cyclin D1^LP^-CDK6 complexes had 53.8±2.4-fold higher activity towards the helix-docking substrate than the non-docking substrate, which was similar to WT cyclin D1-CDK6 complexes indicating that the LP-docking site was not responsible for helix-based docking (**Fig. 1C-D**). Taken together, these data indicate that none of the previously identified G1 cyclin docking sites are directly responsible for helix-based docking.

### Identification of cyclin D1’s helix-docking site using synthetic substrates

Since mutation of known cyclin docking sites did not identify the helix-docking site, we sought to mutate additional regions of cyclin D1. Cyclin D has two lobes, known as cyclin box folds, each comprising five alpha helices^5,7,29^. The N-terminal cyclin box fold binds CDK4 or CDK6 and contains the A1-A5 alpha helices. Notably, the A1 helix contains the hydrophobic patch that binds RxL and related motifs^37–39^. The C-terminal cyclin box fold contains the A1’-A5’ alpha helices, which have not previously been reported to control substrate recognition. The remaining N-terminus is mostly disordered with a single alpha helix preceding the N-terminal cyclin box fold, while most of the amino acids C-terminal to the C-terminal cyclin box fold are also disordered^29^. We therefore generated CDK6 complexes containing thirteen different cyclin D1 variants with mutations in residues on either the unstructured N-terminus, the N-terminal cyclin box fold, the C-terminal cyclin box fold, or the unstructured C-terminus. More specifically, we truncated the first 40 amino acids (ΔN) or the last 25 amino acids (ΔC) and generated eleven individual alpha helix alanine substitution variants (**Fig. 1E**). For each helix variant, we compared structural data and helical predictions for the cyclins D, E, A, and B to identify amino acids unique to cyclin D because only cyclin D exhibits Rb helix-based docking. We reasoned that some residues unique to cyclin D would form a hydrophobic face that docks the Rb C-terminal helix.

To identify the interface on cyclin D1 responsible for helix-based docking, we tested if these cyclin D1 mutations impacted cyclin D1-CDK6’s ability to phosphorylate the same set of synthetic substrates described above. *I.e.*, we want the intrinsic kinase activity of the cyclin D1-CDK6 variant enzyme to be maintained, its RxL-based docking to be maintained, but its helix-based docking to be eliminated. We observed that most of the mutations caused reduction in phosphorylation of the synthetic non-docking CDK substrate reporter, suggesting many of these truncations and alanine substitutions resulted in cyclin D1-CDK6 losing its intrinsic kinase activity (**Fig. 1F-G**). This loss in activity may be caused by destabilization of enzyme structure or decreased CDK binding. The N-terminal cyclin box fold makes contact with CDK, so helix substitution variants likely lost kinase activity through reduced CDK binding. One of the few cyclin D1 variants that demonstrated wild-type kinase activity and RxL-based docking was cyclin D1^A2’^, which had alanine substitutions in the C-terminal cyclin box fold’s A2’ helix. The cyclin D1^A2’^-CDK6 complex exhibited 279.3±12.4-fold higher activity towards the RxL-docking substrate than the non-docking substrate, which was similar to wildtype cyclin D1. Importantly, this variant lost helix-based docking completely and only had 1.4±0.1-fold higher activity towards the helix-docking substrate than the non-docking substrate. This led us to hypothesize that the A2’ helix forms a key part of the interface through which cyclin D1 docks the Rb C-terminal helix (**Fig. 2A**). We note that the variant of cyclin D1 where residues in the A5’ helix were substituted with alanines, cyclin D1^A5’^, phenocopies cyclin D1^LLxxxL^. Since the LLxxxL motif is also within the A5’ helix, this supports a role for the A5’ helix in contributing to Rb C-terminal helix docking by the A2’ helix.

**Figure 2:**
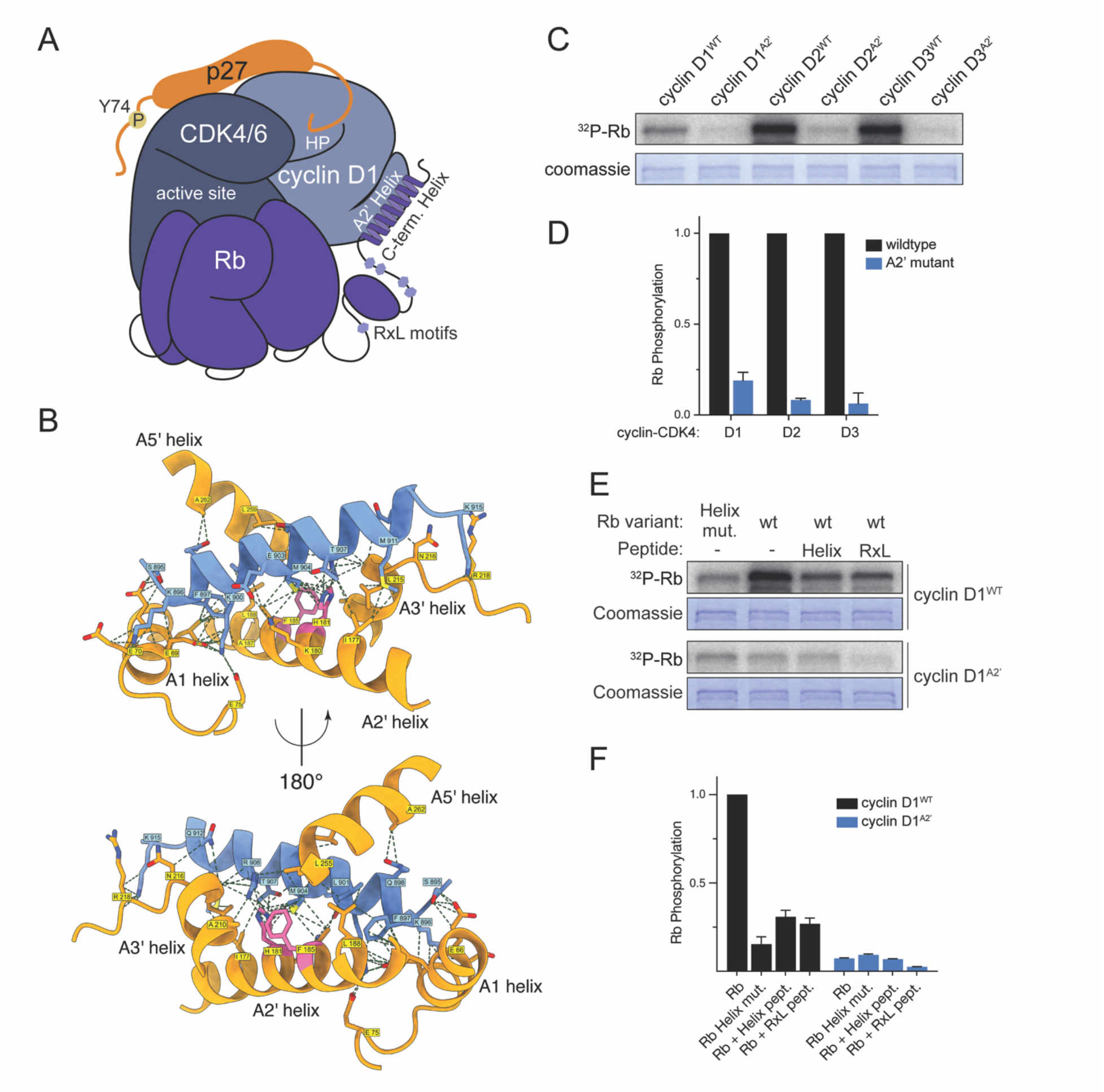
Cyclin D’s A2’ helix docks Rb to target it for phosphorylation. A. Schematic showing docking interactions between the cyclin D-CDK4/6-p27 complex and Rb. The cyclin D A2’ helix docks the Rb C-terminal helix. B. AlphaFold model of the protein complex containing human Cyclin D1, CDK6, and the Rb C-terminal helix (amino acids 895-915). C. Representative *in vitro* kinase assays using the indicated full-length Rb variant with either wild-type cyclin D1, D2, or D3 (*e.g.*, cyclin D1^WT^-CDK6) or the indicated cyclin D A2’ variants lacking helix-based docking. D. Quantification of C (n=3). E. Representative *in vitro* kinase assays using the indicated full-length Rb variant with either wild-type cyclin D1 (cyclin D1^WT^-CDK6) or the cyclin D1 mutant that lacks helix-based docking (cyclinD1^A2’’^-CDK6). Rb^WT^ denotes wild-type Rb and Rb^Helix mut.^ denotes an Rb variant where the helix-based docking interface residues F897, L901, and R908 are substituted with alanines. Where specified, competitive docking peptides were added to each assay. The helix peptide is a peptide derived from 20 amino acids of the Rb C-terminal helix and the RxL peptide is derived from the 9 amino acids of the *S. cerevisiae* CDK inhibitor protein Sic1. F. Quantification of kinase assays from F (n=3).

To further elucidate how the Rb C-terminal helix interacts with Cyclin D1, we used AlphaFold Multimer to model the cyclin D1-CDK6 complex bound to an Rb helix peptide (residues 895–915). The top-ranked multimeric model exhibited high confidence, with uniformly low predicted aligned error (PAE) across the Rb-helix–Cyclin D1 interface (**Fig. S1A-B**). We then mapped residue-level contacts in the best-scoring model and identified four main interaction regions (**Fig. 2B**, Supplementary Table 1 for a list of all contact residues). First, a dominant interface between the Rb helix and the cyclin D1 A2′ helix featured numerous contacts involving cyclin D1 residues I177, K180, H181, T184, F185, A187, and L188. Second, the Rb helix interacted with the cyclin D1 LLxxxL docking region on the A5′ helix, centered on residues L255, L259, and A262. Third, the helix contributed additional contacts, with the N-terminal segment contacting the A1 helix (L65, E66, E69, E70, E75) and the C-terminal segment contacting the A3′ helix (A210, L215, N216, R218). To test this extended docking interface in more detail, we used SIMBA, a yeast-based assay for protein-peptide binding, to measure the RxL and helix docking activity across twelve single or double mutants predicted by AlphaFold^40^. This confirmed that mutations in the A2’ helix (K180A, H181A, T184A, F185A), the A5’ helix (L255A, L259A), and the A1 helix (L65A) can either individually or additively decrease docking to the RBL1 helix (**Fig. S2A**). Collectively, our in silico structure modeling highlights the A2′ helix as the principal binding region for the Rb helix with additional contributions from the A1, A3′ and A5’ helices.

### Deep mutational scanning of cyclin D docking helices reveals binding determinants

To more deeply understand the motif binding determinants predicted by the AlphaFold model, we performed deep mutational scanning of RBL1 and RBL2 peptides and used SIMBA to test binding to cyclins D1, D2, and D3 (**Fig. S2B-C**). In each case, the strongest preferences were observed at three hydrophobic positions followed by a charged C-terminal dipeptide, [DE]R, defining the core motif as ΦxxxΦxxΦxx[DE]R. Each hydrophobic position tolerated multiple aliphatic sidechains, and the first position also permitted aromatic groups (WFY). Intervening positions were less selective, Pro residues were disruptive throughout, and the C-terminal Arg position was the most stringent, as even Lys substitution was strongly disfavored. Notably, Rb contains suboptimal residues at 3 of the 5 core positions (FxxxLxxMxxTR), whereas RBL1 and RBL2 contain the most-favored residues at all 5 positions. While each cyclin showed similar preferences, cyclin D3 tolerated more sequence variation and showed the strongest helix binding (with the rank order D3 > D1 > D2) even when positions were changed to cyclin D1-favored sequences (**Fig. S2D**). To compare the peptide binding requirements with the predicted structural interface, we used the SIMBA data to calculate the Gini coefficient (a measure of inequality) at each peptide position and extracted the fraction of residue surface area from AlphaFold models of multiple cyclin-peptide complexes (**Fig. S2E**). The observed preferences agree well with the predicted peptide interface, and these results collectively define the quantitative binding contributions at each position along the helix.

### The cyclin D1 A2’ helix docks Rb

We next sought to test whether the A2’ helix, identified through assays with synthetic substrates and the AlphaFold model, is required for docking the full-length Rb protein (**Fig. 2A-B**). It is well established that most of the docking interactions on Rb take place C-terminal to the last CDK phosphorylation site T826^24,28,41^. These major docking regions are Rb 830-892, which contains RxL motifs, and Rb 895-915, which contains the C-terminal docking helix. Since we previously found that all human and mouse cyclin Ds use helix-docking to target Rb for phosphorylation^24^, we sought to test if all these cyclin Ds use the A2’ helix to achieve this. We aligned previously published cyclin-CDK complex structures and compared the A2’ helix sequences of cyclin Ds with the other cell cycle cyclins E, A, and B and determined that only cyclins D2 and D3 have conserved interface residues with the cyclin D1 A2’ helix (**Fig. S3A-C**). To test if the alanine substitutions in A2’ helix in cyclin D2 and D3 also disrupt helix-based docking, we performed *in vitro* kinase assays with the full length Rb and compared the wild type with A2’ helix mutant versions of cyclin D2 and D3. Mutation of the A2’ helices in cyclins D1, D2, and D3 had similar effects and reduced the ability of these kinase complexes to phosphorylate Rb by 5.3±0.1-fold, 12.0±0.06-fold, and 15.9±0.5-fold, respectively, compared to wild-type (**Fig. 2C-D**).

To test if the newly discovered cyclin D1^A2’^ variant was also unable to use helix-based docking to target the full-length Rb protein, we performed *in vitro* kinase assays using wild-type Rb (Rb^WT^) and an Rb variant with alanine substitutions at three of the core docking-helix residues F897, L901, and R908 (Rb^Helix^ ^mut.^)^24^. The cyclin D1^A2’^ and Rb^Helix^ ^mut.^ variants had similar effects and reduced Rb phosphorylation by 13.7±0.05-fold and 6.5±0.3-fold, respectively (**Fig. 2E-F**). When we combined these two different mutations in one assay, by phosphorylating Rb^Helix^ ^mut.^ with cyclin D1^A2’^-CDK6, we observed no additional loss of phosphorylation (10.6±0.04-fold reduction). This is consistent with these two variants having defects in the same docking interaction. To further test the role of the A2’ helix in Rb docking, we performed *in vitro* kinase assays with cyclin D1^WT^ or cyclin D1^A2’^ and with peptides containing either RxL or Rb C-terminal helix sequences, which reversibly inhibit RxL-based or helix-based docking, respectively. As expected, the RxL peptide further reduced Rb phosphorylation by cyclin D1^A2’^-CDK6 to background levels, while the Rb C-terminal helix peptide was unable to further inhibit cyclin D1^A2’^-CDK6 (**Fig. 2E-F**). Taken together, these data argue that the A2’ helix of the cyclin D family is required for helix-based docking of the full-length Rb protein and is not involved in RxL-based docking.

### Mutation of cyclin D1’s A2’ helix reduces proliferation and increases cell size

Having identified mutations in the A2’ helix of cyclin D1 that disrupted helix-based docking *in vitro*, we aimed to test if these mutations affected the G1/S transition in cells. To do this, we generated RPE-1 cell lines that contain a doxycycline-inducible cyclin D1 replacement system. This system allows conditional expression of an shRNA targeting endogenous cyclin D1 and an exogenous, shRNA-resistant, FLAG-tagged cyclin D1 (**Fig. 3A**)^16^. We note that the RPE-1 cell line is a non-transformed, hTERT1-immortalized cell line that mainly expresses cyclin D1 out of the D-type cyclins. Using this replacement system, we introduced cyclin D1^WT^, cyclin D1^HP^, or two different cyclin D1^A2’^ variants into RPE-1 cells while knocking down the endogenous cyclin D1. As expected, shRNA knockdown of cyclin D1 alone reduced Rb phosphorylation and arrested cells in G1, while induction of shRNA-resistant cyclin D1^WT^ above endogenous levels increased Rb phosphorylation and reduced the fraction of cells in G1 from 71.4±2.9% to 35.2±1.6% (**Fig. 3B-C**). Compared to cyclin D1^WT^, expression of either of the two different helix-based docking deficient cyclin D variants was less effective in promoting Rb phosphorylation and cell proliferation (52.1±1.8 or 47.4±1.5% cells in G1 compared to 35.2±1.6% for the expression of cyclin D1^WT^). In addition, the expression of WT cyclin D1 resulted in smaller cells than the expression of cyclin D1^A2’^ as is expected if this mutation slowed down cell cycle progression (**Fig. 3D**). We note that the replacement system we used leads to overexpression of cyclin D1 compared to endogenous levels. Taken together, our results suggest mutation of the cyclin D A2’ helix inhibits the ability of cyclin D to drive the G1/S transition in a similar way to the mutation of the Rb C-terminal helix^24^.

**Figure 3:**
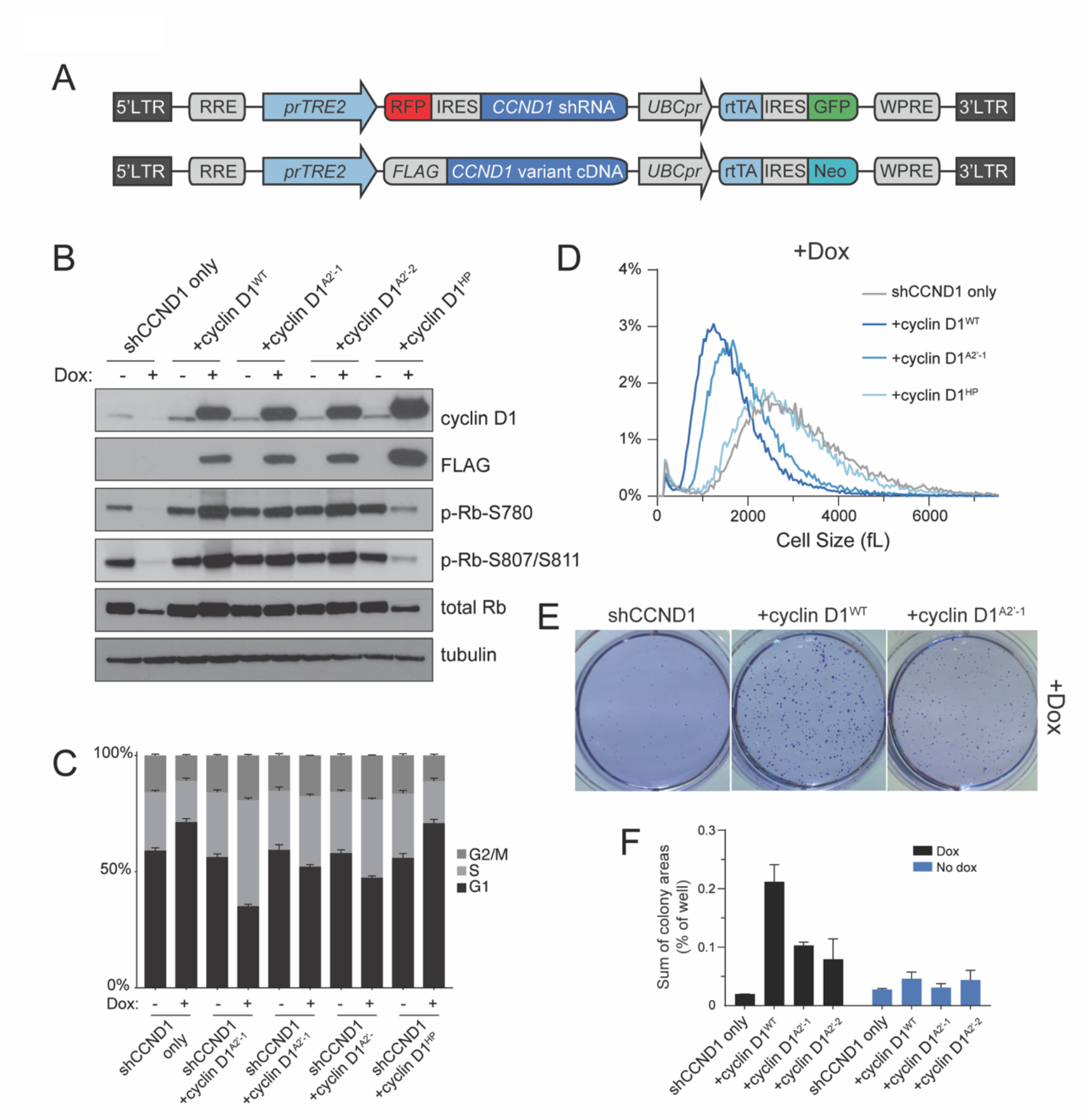
Mutation of cyclin D1’s A2’ helix reduces proliferation and increases cell size. A. Lentiviral integration constructs containing doxycycline-inducible shRNA targeting the 3’ UTR of *CCND1* and doxycycline-inducible *CCND1* cDNA fused to a FLAG affinity tag sequence. Constructs were integrated into RPE-1 cells. B. Immunoblot analysis of lysates from RPE-1 cells containing the constructs indicated in (A) that were treated with doxycycline for 48 hours to knockdown the endogenous cyclin D1 and express the indicated *CCND1* mutant. WT denotes wild-type cyclin D1, A2’-1 and A2’-2 denote two different sets of alanine mutations in cyclin D1’s A2’ helix, and HP denotes mutation of the hydrophobic patch. C. Flow cytometry analysis of cell cycle phase distribution using EdU incorporation and DAPI staining after 48 hours of doxycycline treatment as in B. D. Cell size analysis using a Coulter counter of the indicated RPE-1 cell lines after 48 hours of doxycycline treatment as in B. E. Representative images of the indicated RPE-1 cell lines growing on soft agar in media containing doxycycline. F. Quantification of colony area for soft agar growth assays in E.

Notably, we found that expression of the cyclin D1^HP^ variant in cells caused no changes to Rb phosphorylation, G1 fraction, and cell size compared to cyclin D1 knockdown alone (**Fig. 3B-D**), suggesting that the cyclin D1^HP^ variant is either entirely unable to bind Rb or properly form an active cyclin D1^HP^-CDK-p27 trimer complex^7^. Deeper investigation into this activation mechanism and the role of p27 in helix-based docking is required to distinguish between these possibilities.

We next sought to further test the function of the cyclin D A2’ helix in promoting proliferation. To do this we examined the effect on colony formation in soft agar^42^ in cells expressing different cyclin D1 variants. Indeed, proliferation on soft agar was enhanced by the expression of cyclin D1^WT^ more than by the expression of cyclin D1^A2’^, consistent with our model (**Fig. 3E-F**).

### The cyclin D-CDK4-p27 trimer is assembled through HP docking

That the mutation of cyclin D’s hydrophobic patch had a phenotype similar to cyclin D knockdown suggested that this mutant failed to assemble active cyclin D-CDK4/6 complexes (**Fig. 4A**). Consistent with this model, it was previously shown that cyclin D’s hydrophobic patch binds p27, which can serve as a kinase complex assembly factor^7,31^. To test if the cyclin D hydrophobic patch is required to properly form an active cyclin D-CDK4/6 complex in cells via the p27 assembly factor we used our cyclin D1 replacement system described above to knockdown the endogenous cyclin D1 and re-express different cyclin D1 variant proteins fused to a FLAG epitope tag. p27 was pulled down by FLAG-cyclin D1^WT^ and detected by immunoblot but was not pulled down by FLAG-cyclin D1^HP^ (**Fig. 4B**). In contrast, p27 was pulled down by cyclin D1 variant proteins with mutations to the A2’ helix. This is consistent with the cyclin D-CDK4/6 assembly factor operating through cyclin D’s hydrophobic patch, but not the A2’ helix.

**Figure 4:**
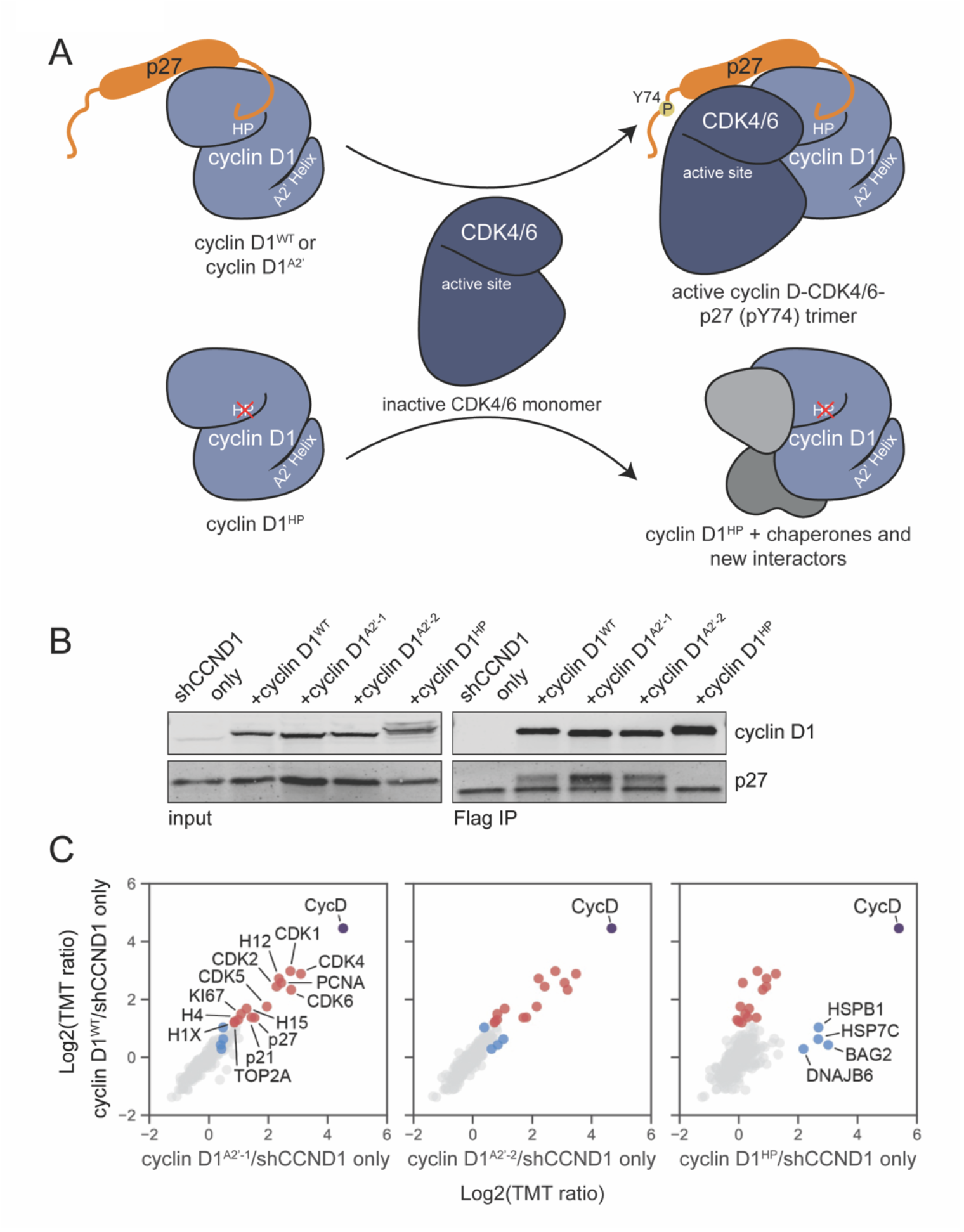
Cyclin D’s hydrophobic patch, not the A2’ helix, is required for cyclin D-CDK4-p27 trimer assembly. A. Model schematic showing cyclin D-CDK4-p27 trimer assembly uses cyclin D’s hydrophobic patch (HP) not the A2’ helix docking site. B. Immunoblots performed after immunoprecipitation of the indicated FLAG-cyclin D protein from lysates of RPE-1 cells expressing the indicated cyclin D1 variant protein as described in Fig. 2A-B. C. Mass spectrometry analysis of samples collected from immunoprecipitation of the indicated FLAG-cyclin D protein as in B (see methods). A total of 293 proteins were detected (>2 unique peptides). Proteins enriched in FLAG-cyclin D1^WT^ are labeled in red and proteins enriched in FLAG-cyclin D1^HP^ are labeled in blue. Bait protein (FLAG-cyclin D1) is labeled in purple.

To more comprehensively test how cyclin D’s interfaces are responsible for its protein interactions, we performed a series of cyclin D1-FLAG pulldown experiments, as described above, followed by mass spectrometry (see Methods). Mutation of the A2’ helix did not appreciably affect the set of proteins pulled down by cyclin D1, which still included the expected CDK partner proteins and p27 (**Fig. 4B-C**). In contrast, mutation of the hydrophobic patch reduced this set of cyclin D1-interacting proteins to background or near background levels, while a few new interactions appeared, including chaperone proteins (**Fig. 4C**). This supports the model that HP docking is important for trimer assembly and that the impact of cyclin D^A2’^ on Rb phosphorylation and the G1/S transition is through disrupting the cyclin D-Rb docking interaction.

### Cyclin D docks Rb’s C-terminus when it is in a helical conformation

We previously showed that Rb’s C-terminus docks cyclin D when it is in a helical conformation as proline substitutions disrupting the helical conformation reduce the ability of cyclin D-CDK4/6 complexes to phosphorylate Rb^24^. The Rb C-terminal helix was found to be in the helical conformation ∼30% of the time, so it should be possible to enhance the helicity of Rb’s C-terminal docking region to improve cyclin D docking. To increase the helicity of Rb helix peptides, we used hydrocarbon stapling to covalently constrain the peptide into a helical conformation at multiple positions throughout the peptide (**Fig. 5A**). We cross-linked residues within the peptide that were 3 residues apart and did not modify any of the interface residues involved in docking. We tested these stapled peptides for increased docking activity compared to the unstapled Rb peptide using *in vitro* kinase assays of Rb with cyclin D1-CDK6 as described above. Four of the seven stapled peptides had improved docking activity over the unstapled helix peptide (**Fig. S4A-B**). Three of these improved docking peptides, including the strongest inhibitor, Stapled-1, were stapled near the N-terminus of the peptide. The staples placed more centrally in the peptide did not improve inhibition. The IC_50_ of Stapled-1 is 0.64 µM (95% CI: 0.36-1.12 μM), a 26-fold improvement over the unstapled Rb helix peptide’s IC_50_ of 16.72 μM (95% CI: 10.27-29.74 μM) (**Fig. 5B**). When we mutated the core helix-docking residues F897, L901, and R908 of the Rb Helix peptide to alanine (Rb Helix 3A stapled peptide), Rb phosphorylation was no longer inhibited (**Fig. 5B**). Taken together, these peptide experiments show that enhancing the helicity of the Rb C-terminal helix improves its ability to dock cyclin D.

**Figure 5:**
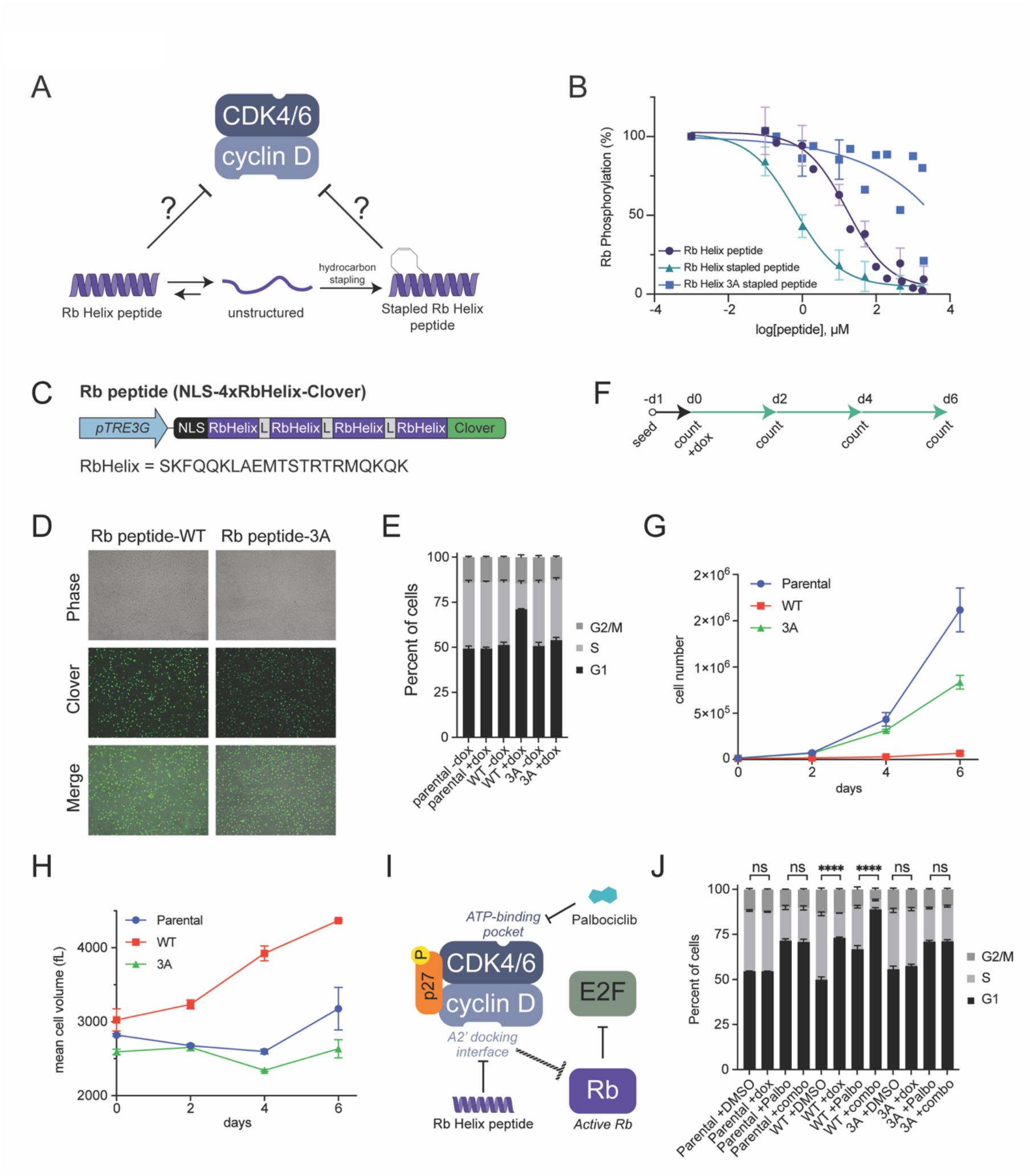
The Rb C-terminal helix peptide inhibits cyclin D-CDK4 and cell proliferation. A. Schematic showing how Rb(895-915)-derived peptides adopt both unstructured and helical conformations and how hydrocarbon stapling can force peptides into a helical conformation. B. Quantification of *in vitro* kinase assays using cyclin D1-CDK6 and full length Rb proteins. The reaction also contained the indicated concentration of peptides derived from Rb’s C-terminal helix. 3A denotes a peptide in which the 3 interface residues were substituted with alanines. Stapled peptide indicates a peptide with an N-terminal hydrocarbon staple that had the strongest inhibition of CDK6 activity in Fig. S4A-B. C. PiggyBac integration constructs containing doxycycline-inducible Rb peptide (NLS-4xRbHelix-Clover). Constructs were integrated into RPE-1 cells. D. Representative phase contrast and fluorescent images of cells expressing WT or 3A Rb peptide after 48 hours of doxycycline treatment. E. Flow cytometry analysis of cell cycle phase distribution using EdU incorporation and DAPI staining of control RPE-1 cells (parental) or RPE-1 cells expressing WT or 3A Rb peptide after 48 hours of doxycycline treatment as in D. n = 4. F. Schematic of timecourse experiment measuring cell numbers and size over six days of doxycycline treatment for control RPE-1 cells (parental) or RPE-1 cells expressing WT or 3A Rb peptide. G. Cell number analysis using a Coulter counter of cell lines as described in F. n = 3. H. Cell size analysis using a Coulter counter of cell lines as described in F. n = 3. I. Schematic of combined cyclin D inhibition by a helix peptide derived from the Rb C-terminus and the CDK4/6i inhibitor Palbociclib. J. Flow cytometry analysis of cell cycle phase distribution using EdU incorporation and DAPI staining of control RPE-1 cells (parental) or RPE-1 cells expressing WT or 3A Rb peptide after 48 hours of DMSO, doxycycline (dox), 50 nM Palbociclib (Palbo), or combined doxycycline and 50 nM Palbociclib (combo). n = 3. ns (p > 0.05), **** (p ≤ 0.0001).

### A cyclin D docking inhibitor is more potent than small molecule CDK4 inhibitors at targeting cyclin D-CDK4/6-p27 trimer complexes

While Rb-helix-based peptides can serve as docking inhibitors of cyclin D-CDK4,6 dimer complexes, much if not most of the cyclin D-CDK4/6 complexes in cells may exist as a trimer with p21 or p27, which could have distinct biochemical properties (**Fig. S4C**). While p21 and p27 are normally thought of as CDK2 inhibitors, they can also activate or inhibit cyclin D-CDK4/6 complexes. The activating or inhibitory roles of p21 and p27 depend on their phosphorylation^32^. It is clear p21 and p27 serve as assembly factors because mouse fibroblasts lacking both proteins failed to assemble cyclin D-CDK4 complexes^31^, and active CDK4 complexes contain phosphorylated p27 that alters its ATP binding site^7^. To compare the biochemical activities of the dimer and trimer cyclin D-CDK4 complexes, we first performed a series of *in vitro* kinase assays with full length Rb protein. The cyclin D1-CDK4 dimer is more active than the cyclin D1-CDK4-p27 trimer, but both complexes were specific for Rb (**Fig. S4D-E**).

Interestingly, a recent report found that inhibitors targeting CDK4’s ATP binding pocket, such as palbociclib, were poor inhibitors of cyclin D-CDK4-p27 trimers and proposed that these molecules inhibited the G1/S transition by binding CDK4/6 monomers to prevent the assembly of cyclin D-CDK4/6 active complexes^7^. Consistent with this report, we found that palbociclib was a poor inhibitor of cyclin D1-CDK4-p27 trimers **(Fig. S4F)**. In contrast to ATP competitors targeting CDK4, our peptides target the docking site on cyclin D to inhibit the ability of cyclin D-CDK4 to phosphorylate Rb (**Fig. 5A-B, S4F**). This suggests that such docking inhibitors targeting cyclin D may be better suited to inhibit our cyclin D1-CDK4-p27 trimers. To test this, we performed *in vitro* kinase reactions with the stapled Rb-helix peptide inhibitor described above to measure the inhibition constant (K_i_) for the cyclin D1-CDK4-p27 phosphorylation of Rb. This allowed us to measure the IC_50_ with the Stapled-1 peptide as 19.8 µM (**Fig. S4G**). In contrast, the Stapled-1 peptide with substituted interface residues (3A stapled peptide) was unable to inhibit the ability of the trimer to phosphorylate Rb. This shows that the cyclin D1-CDK4-p27 trimer uses the helical docking mechanism to target Rb for phosphorylation and that the hydrophobic patch is used for p21/p27 based assembly^7^. Taken together, our experiments show that helical peptides targeting cyclin D docking may be more versatile inhibitors of cyclin D-CDK4/6 activity since they would be able to effectively inhibit both dimer and trimer complexes.

### A cyclin D docking inhibitor can target non-canonical cyclin D-CDK complexes

During normal cell cycles, D-type cyclins form canonical complexes with CDK4 and CDK6. This regulation is frequently disrupted in cancer, where overexpression of cyclin D leads to the formation of non-canonical complexes with other Cdks, such as CDK1, CDK2, and CDK5 (**Fig. 3C)**^43^. Because CDK4/6 inhibitors such as palbociclib were developed to target cyclin D-CDK4/6 complexes, we hypothesized that these non-canonical complexes, like cyclin D1-CDK2, would be insensitive to palbociclib inhibition and could therefore contribute to drug resistance. To test this hypothesis, we performed in vitro kinase assays measuring phosphorylation of the model substrate RbC, which is derived from an Rb fragment (772-928) that contains both phosphorylation sites and intact helix-based and RxL-based docking sequences^44,45^. We examined phosphorylation of this fragment by cyclin D1-CDK6 or cyclin D1-CDK2 complexes in the presence of increasing concentrations of palbociclib. As expected, palbociclib potently inhibited RbC phosphorylation by cyclin D1-CDK6, with an IC_50_ of approximately 6 nM. In contrast, cyclin D1-CDK2 complexes were largely insensitive to palbociclib inhibition (**Fig. S4H**). Since both canonical cyclin D1-CDK4/6 and non-canonical cyclin D1-Cdk2 complexes should rely on a helix-based docking mechanism to phosphorylate Rb, we hypothesized that both types of complexes would be sensitive to inhibition by a helix-based docking inhibitor derived from the Rb C-terminus. To test whether the A2′ helix docking inhibitor can target both types of cyclin D complexes, we performed in vitro kinase assays using increasing concentrations of the stapled Rb helix peptide. The stapled peptide blocked Rb phosphorylation by cyclin D1-CDK2 with an IC_50_ of approximately 250 nM, comparable to the IC_50_ observed for the canonical cyclin D1-CDK6 complex (**Fig. S4I**). Taken together, these experiments suggest that molecules targeting cyclin D-based docking can be potent inhibitors of cell proliferation in scenarios where CDK4/6 inhibitors are not effective.

### The Rb C-terminal helix peptide reduces cell proliferation and increases cell size

Having established that targeting cyclin D’s A2’ helix with the Rb-helix-based peptide potently inhibits all forms of cyclin D-CDK activity in kinase assays, we next wanted to target the cyclin D interface in a cellular context. Inhibitory peptides targeting protein-protein interactions are a promising therapeutic strategy, as they offer high specificity and can target broad protein interfaces^46^. To target cyclin D’s A2’ helix in cells, we chose to express multiple linked copies of the Rb helix, since previous work has shown that linking multiple copies of the same binding motif with a long, flexible linker can improve avidity and affinity for a target^48^. As a result, we stably integrated an inducible construct to express four Rb helices linked by a flexible GS-linker, tagged with an N-terminal NLS and a C-terminal Clover fluorescent protein (Rb peptide; NLS-4xRbHelix-Clover; **Fig. 5C-D**). Induction of Rb peptide increased the fraction of cells in G1 from 51.3±1.5% without doxycycline to 71.1±0.3% with doxycycline induction (**Fig. 5E**). In contrast, cell cycle progression was the same in cells expressing a variant in which the three docking interface residues were substituted with alanines (Rb peptide-3A; NLS-4xRbHelix-3A-Clover; G1: 50.7±2.1% to 53.4±1.5%) or an unmodified parental RPE-1 control (G1: 49.3±1.3% to 49.3±0.8%). We also generated untagged, HA-tagged, and T2A-Clover cleavage variants of Rb peptide that similarly increased the percent of cells in G1 after doxycycline treatment (**Fig. S5A-B**). In addition, the expression of Rb peptide reduced cell numbers, increased cell size, and caused a reduction in Rb phosphorylation and overall Rb amount (**Fig. 5F-H, S5C**), which is consistent with inhibition of cyclin D-CDK4/6 complexes^19,24^. These results align with work showing that RBL2 peptide expression affects cancer cell proliferation^47^ and, together, demonstrate the potential of targeting the cyclin D subunit to stop cell proliferation.

### The wildtype Rb C-terminal helix peptide specifically inhibits proliferation in cells

While these results suggest that Rb peptide is a potent peptide inhibitor of cyclin D-CDK4/6, we observed that the WT variant of the peptide is expressed at a higher level than the 3A variant (**Fig. 5D, S5C**). To rule out any effects caused by differences in expression level, we induced Rb peptide expression with a range of doxycycline concentrations between 0.01 µg/mL and 2 µg/mL. Only the WT, but not the 3A, variant of Rb peptide exhibited a dose-dependent response in the percent of cells in G1 (**Fig. S5D-E**). Furthermore, based on the Clover signal, we also analyzed cell cycle phase distribution of cells divided into four equal bins of Rb peptide expression (**Fig. S5F**). We observed that at no, or relatively low levels of Clover expression, cells expressing WT or 3A were indistinguishable. However, at all higher levels of Clover expression, the percent of WT-expressing cells in G1 was significantly increased compared to 3A-expressing cells (**Fig. S5G-H**).

### Helix peptide expression improves the effect of CDK4/6i treatment

We next sought to test whether combined targeting of both the ATP-binding pocket and the A2’ docking interface of cyclin D-CDK4/6 would improve upon either individual inhibition strategy (**Fig. 5I**). To do this, we analyzed cell cycle phase distribution in cells treated with DMSO, Rb peptide alone (doxycycline to induce NLS-4xRbHelix-Clover), palbociclib alone, or a combination of Rb peptide and palbociclib. While palbociclib alone (G1: 71.5±1.1%) and induction of Rb peptide alone (G1: 73.1±0.4%) had similar effects, combined treatment increased the percentage of G1 cells to 88.9±1.0% (**Fig. 5J**). Thus, these cell-based peptide inhibitor experiments reveal an alternative and promising approach for cyclin D-CDK4/6 complex inhibition, independent of the ATP-binding pocket of CDK4/6.

## Discussion

The eukaryotic cell division cycle entails a sequence of events starting with transcriptional activation through to DNA replication, mitosis, and cytokinesis. The timing of these events is in part regulated by the phosphorylation of key substrates by cyclin-dependent kinases whose aggregate activity increases monotonically through the cell cycle^49–52^. Consistent with this picture, G1 cyclin-CDK complexes have a lower activity towards a small peptide derived from histone H1 containing CDK consensus phosphorylation sites compared to the S phase and mitotic cyclin-CDK complexes that are assembled later in the cell cycle^53^. While this model first emerged from yeast studies, we recently confirmed a similar progressive increase in intrinsic kinase activity of animal cyclin-CDK complexes through the cell cycle, where cyclin D-CDK4/6 complexes exhibited the lowest intrinsic H1 kinase activity^24,54^. Similarly, the earliest activated budding yeast cyclin-CDK complex, Cln3-CDK1, has extremely low intrinsic activity^55^. Thus, the low intrinsic activity of the early cyclin-CDK complexes prevents them from triggering events that should take place later in the cell division cycle.

Because of their low intrinsic activity, early cyclin-CDK complexes require specific docking mechanisms to sufficiently phosphorylate their substrates^37^. Docking mechanisms typically confer substrate specificity to cyclin-CDK complexes via the structured cyclin surface binding a short linear motif in an unstructured region of the target protein. The earliest discovered such docking mechanism was a hydrophobic patch on the surface of both yeast and animal cyclins that recognized RxL sequences to phosphorylate substrates initiating DNA replication^28,56^. This same hydrophobic patch region on different cyclins exhibits differential specificity to a variety of short linear motifs. For example, the mitotic cyclin Clb2 prefers LxF motifs, while Clb3 prefers PxxPxF motifs^38,39^. Similarly, while the hydrophobic patches of human cyclins E and A target RxL sequences, a recent comparison of cyclin E, A, and B revealed distinct specificities for docking sequences^28,39,57^. Thus, although there is some overlap in specificity, such as RxL recognition, ostensibly similar hydrophobic patches can dock distinct motifs on target proteins. Moreover, cyclin docking interactions are not limited to the hydrophobic patch as the budding yeast G1 cyclins use a different groove to recognize an LP (or LLPP) motif^36,53^.

Like other G1 cyclin-CDK complexes, cyclin D-CDK4/6 complexes need to compensate for their low intrinsic activity by using a highly specific mechanism to dock a helix within the largely unstructured region in Rb’s C-terminus^24^. This cyclin D-Rb docking interaction promotes Rb phosphorylation and progression through the G1/S transition. Here, we report the identification of a region on cyclin D’s surface, the A2’ helix, likely responsible for docking Rb. Expression of a cyclin D variant containing mutations of amino acids within the A2’ helix decreased Rb phosphorylation and slowed G1/S progression. Interestingly, the A2’ helix on cyclin D is not any of the previously identified protein binding surfaces of cyclin D including the hydrophobic patch, the LP-binding groove, the LxxLL motif, and the LxCxE motif^23,26,27,34–36^. Rather, the A2’ helix represents a novel substrate docking site on all three human cyclin D proteins that is crucial for driving the G1/S transition.

Our identification and mapping of the cyclin D-Rb docking interaction provides an additional target for cancer drugs because deregulation of the G1/S transition and the cyclin D-CDK4/6-Rb pathway is a hallmark of many cancer cells^1–3^. In addition, targeting cyclin D’s A2’ docking interface provides an opportunity to inhibit the trimer and non-canonical forms of cyclin D complexes (**Fig. 6**). Currently, small molecule cancer drugs inhibiting the G1/S transition target the ATP binding pocket of CDK4 and CDK6^3,58,59^. Although this crop of small molecule CDK4/6 ATP pocket inhibitors have been observed to limit disease progression, they have modest if any effects on overall survival, suggesting there is significant scope to improve on current therapies^60–62^. One reason for this could be that while ATP pocket inhibitors can inhibit cyclin D-CDK4/6 dimer complexes, they cannot inhibit cyclin D-CDK4/6 trimer complexes including the assembly factors p27 or p21^7^. Importantly, it is possible that these trimers may be the primary active complexes in some cells because MEFs lacking the assembly factors p21 and p27 failed to form active CDK4 complexes^6,31^. In addition, CDK4/6 inhibitors are unable to target non-canonical cyclin D complexes, such as those with CDK2. In contrast to the CDK4 ATP pocket inhibitor palbociclib, Rb-derived helix peptides can be used as a cyclin D docking site inhibitor to prevent cyclin D-CDK4-p27 trimers and non-canonical cyclin D-CDK2 complexes from phosphorylating Rb and thereby inhibiting cell cycle progression. These proof-of-principle experiments show how new drugs targeting cyclin D substrate recognition might improve on current cancer therapy by targeting all forms of active cyclin D-CDK4/6 complexes.

**Figure 6:**
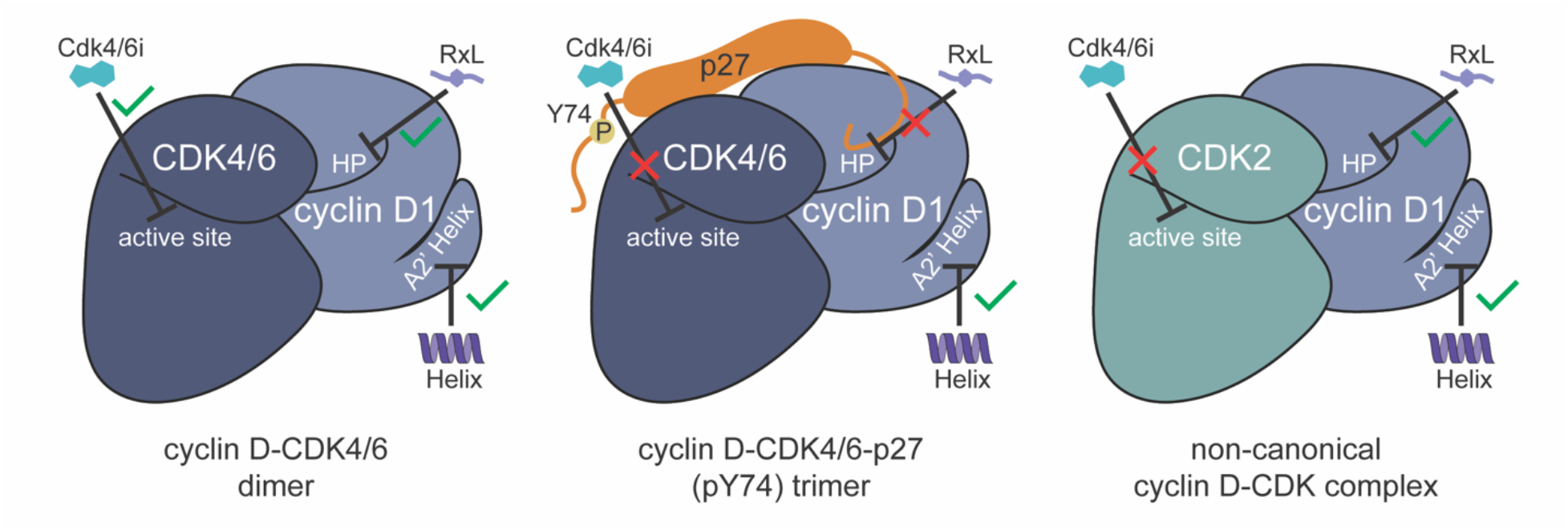
Model of the cyclin D A2’ helix as a novel target for therapeutics to inhibit cell proliferation. Unlike conventional active site inhibition with Cdk4/6i or targeting the HP-RxL interaction, A2’ helix docking inhibition is possible for dimer, trimer, and non-canonical forms of cyclin D complexes.

## Methods

### Cell Culture

Non-transformed human retina epithelial cells RPE-1 cells transformed with hTERT were cultured in Dulbecco’s Modified Eagle’s Medium (DMEM) supplemented with 5% Fetal Bovine Serum (FBS) and antibiotics (100 units/mL Penicillin and 100 μg/mL Streptomycin).

### Doxycycline-inducible cyclin D replacement system and cell line construction

The pINDUCER system was used to generate doxycycline-inducible *CCND1* shRNA and shRNA-resistant *CCND1* RPE-1 cells^63^. FLAG-tagged *CCND1* was amplified by PCR and was transferred to the pENTR/D-TOPO cloning vector, in accordance with manufacturer’s instructions (Invitrogen, Cat. No. 45-0218). To generate the cyclin variants, we performed site-directed mutagenesis using the QuickChange II XL Site-Directed Mutagenesis Kit (Agilent, Cat. No. 200521). *CCND1* cDNAs were subsequently transferred into pINDUCER20 (Addgene; 44012) with the LR clonase reaction (Invitrogen, cat. No. 11791-020). shRNA that targets the 3’-UTR of *CCND1* mRNA was subcloned in XhoI – MluI sites in pINDUCER11 (Addgene; 44363), as it was described before^63^.

pINDUCER11 and pINDUCER20 lentiviral constructs were packaged in 293T cells by transient transfection, in combination with the envelope plasmid pCMV-VSV-G (Addgene; 8454) and the packaging plasmid pCMV-dR8.2 dvpr (Addgene; 8455). Transfections were carried out using X-tremeGENE 9 DNA Transfection Reagent (Sigma, Cat. No. 6365787001). RPE-1 cells were infected with the lentiviral particles in the presence of 5 mg/mL polybrene (Sigma, Cat. No. 107689). Depending on the selection marker, stable cell lines were selected for expression of the green fluorescence protein by fluorescence-activated cell sorting or resistance to 400 mg/mL G-418 (GIBCO, Cat. No. 10131035). shRNA-mediated knockdown of endogenous cyclin D and expression of Cyclin D variants in stable cell lines were confirmed via immunoblot analysis of cell lysates after 48 hours of 500 ng/mL doxycycline induction.

### Doxycycline-inducible PiggyBac integration vector and cell line construction

gBlocks for NLS-4xRbHelix WT and 3A were synthesized by Integrated DNA Technologies (IDT) and used as templates for PCR. The peptide sequence comprises four Rb C-terminal Helix (Rb 895-915: SKFQQKLAEMTSTRTRMQKQK) linked with three G_4_S glycine-serine linkers (GGGGS GGGGS GGGGS) preceded by an SV40 NLS sequence (PKKKRKV). Gibson assembly was used clone NLS-4xRbH into a PiggyBac integration plasmid, driven by the TRE3G doxycycline-inducible promoter. The NLS-4xRbH-Clover, HA-NLS-4xRbH, and HA-NLS-4xRbH-Clover variants were generated by modifying PCR primers to add HA and T2A sequences and/or using a PiggyBac integration plasmid that contained an in-frame Clover fluorescent protein sequence. Zeocin (final concentration 400 μg/mL; Invivogen ant-zn-05) selection began two days after transfection. Zeocin resistant cells were maintained as polyclonal cell lines and expression was induced with doxycycline (500 ng/mL) for two days or as specified in the experiment.

### Immunoblot analysis

Cells were washed with phosphate-buffered saline (PBS) and total cell lysates were collected in 1X RIPA buffer supplemented with protease (Roche, Cat. No. 04693159001) and phosphatase (Roche, Cat. No. 04906837001) inhibitor-cocktails. Lysates were clarified by centrifugation at 16,000 x g for 10 min at 4°C, separated on Criterion TGX gels (BioRad, Cat. No. 5671084) and transferred onto PVDF membranes (BioRad, Cat. No. 1704273).

Membranes were incubated with the following antibodies: Phospho-Rb (Ser807/811) (D20B12) XP® Rabbit mAb (Cell Signaling Technology #8516), Phospho-Rb (Ser780) (D59B7) Rabbit mAb (Cell Signaling Technology #8180), Rb (4H1) Mouse mAb (Cell Signaling Technology #9309), Monoclonal Anti-α-Tubulin antibody produced in mouse (Sigma-Aldrich T9026), and Rabbit polyclonal cyclin D1 Antibody H-295 (Santa Cruz Biotechnology sc-753).

### Cell size measurements

Cells were washed with PBS, harvested by trypsinization (Gibco 25200056), and resuspended ISOTON II Electrolyte Diluent (Beckman Coulter 8546719). Cell size distributions were measured using a Z2 Coulter Counter (Beckman Coulter).

### Flow Cytometry Analysis

Flow cytometry analysis was performed on Attune NxT Thermo Fisher Scientific cytometer. Cells were prepared for flow cytometry by washing with PBS, trypsinizing, and resuspending PBS. For measurement of newly synthesized DNA by nucleoside analog incorporation, cells were incubated with 10 µM EdU for 30 minutes at 37°C, fixed with 3% formaldehyde in PBS for 10 minutes at 37°C, and permeabilized with 90% methanol for 30 minutes on ice. EdU incorporation was detected using the Click-iT EdU Alexa Fluor 647 Flow Cytometry Assay Kit (Invitrogen C10424) and cells were stained with 3 uM DAPI (Invitrogen D1306). For experiments where GFP expression was also analyzed, EdU incorporation was detected using the Click-iT Plus EdU Alexa Fluor 647 Flow Cytometry Assay Kit (Invitrogen C10634). For cell cycle analysis, we measured EdU signal as a readout for S phase cell population and DAPI fluorescence as a readout for DNA content.

### Soft Agar colony formation assay

40 mL of prewarmed DMEM-F12 (Corning 10-013-CV) medium containing 10% FBS and 5% Penicillin-Streptomycin was mixed with 10 mL 3% SeaPlaque low-melting temperature agarose (Lonza 50101) solution and transferred to 6-well plates to form a 0.6% agarose bottom layer. Plates were incubated at room temperature for 1 hour and 4°C for 10 minutes to set. Cell suspensions of each were prepared by adding 2.5×10^4^ cells to 2.5 mL DMEM-F12 media. 2.5 mL of 0.6% agarose solution was added to the cell suspension for a final top layer agarose concentration of 0.3%. Aliquots of cell suspensions were plated on the bottom layer. The top layers were allowed to harden in the hood for 1 hour and then transferred to 4°C for 15 minutes to solidify. The 6-well plates were then transferred to a 37°C incubator for 3 weeks to observe colony formation. Plates were fed with media either containing 500 ng/ml doxycycline or DMSO as needed to prevent drying. Colonies were stained with 0.1% crystal violet (Fisher Chemical C581-25) for 20 minutes and then washed with water until colonies could be observed. Plates were scanned with a flatbed scanner.

6 well plate images were processed using ImageJ. Scanned 6 well plate images were converted into grayscale cropped well images and between 3-5 regions of interest were selected from each well image for analysis. Grayscale images were processed by manually determining the optimal number of sigmas away from the mean pixel value of each image to distinguish the background and foreground intensity threshold. The intensity threshold was used to generate masks of the images from which colony areas and counts could be extrapolated. Total colony density of the wells of each condition was calculated from the total number of colonies detected from the ROI masks corresponding to each condition normalized to the ROI area.

### FLAG-Cyclin D Immunoprecipitation

The cyclin D mutant immunoprecipitation protocol was adapted from a previous protocol^64^ that was optimized to maximize extracting active Rb in complex with its interactors. This protocol uses a higher salt concentration for lysis to disrupt chromatin and includes 10% glycerol to preserve protein integrity. In brief, RPE-1 cell lines expressing *CCND1* shRNA alone or *CCND1* shRNA with WT or docking mutant *CCND1* cDNAs were treated with 0.5 µg/mL dox for 48 hours to induce the depletion of endogenous cyclin D and replacement with exogenous cyclin D. Approximately 2×10^7^ cells were washed twice with cold PBS pelleted and resuspended in 1 mL E1Agl^250^ buffer (50 mM Hepes-KOH pH 7.4, 250 mM NaCl, 0.1% NP-40, 10% glycerol, 1mM PMSF) with Roche protease and phosphatase inhibitors. Lysates were incubated on ice for 10 minutes and then sonicated using a Model 120 Sonic Dismembrator (Fisherbrand FB120110) for 2×15 sec pulses (45 sec interval) and 50% amplitude. Lysates were centrifuged for 10 minutes at 16,000 x g at 4°C to clear cell debris. To precipitate FLAG-tagged cyclin D variants, the supernatants were incubated with anti-FLAG M2 magnetic beads (Sigma M8823) for 2 hours rotating at 4°C. Beads were washed three times with 1 mL E1Agl150 buffer (50 mM Hepes-KOH pH 7.4, 150 mM NaCl, 0.1% NP-40, 10% glycerol, 1mM PMSF) with a lower salt concentration and proteins were eluted twice with elution buffer (100µg/mL FLAG peptide, Sigma F3290, in TBS; 10mM Tris-HCl pH 7.5, 150 mM NaCl) for western blot or once with 6x Laemmli sample buffer for proteomics analysis.

### Mass spectrometry analysis

After washing, IP resin was resuspended in a 50mM Tris-HCL (7.5pH) buffer with 1% SDS. Protein was eluted from the resin by heat (65C) for 15 minutes. After separating elution from the resin, eluted proteins were reduced using 5mM DTT (10 minutes), capped with 10mM Iodoacetamide (10 minutes), and then precipitated using 50:50 ethanol:acetone solution on ice for 30 minutes. Precipitated protein was then pelleted at max speed for 15 minutes, washed once with 50:50 ethanol:acetone solution, and spun down again. Proteins were re-solubilized in 2 M urea, 50 mM Tris-HCl and 150 mM NaCl, and then digested overnight with 2 ug of Trypsin GOLD at 37 °C. Trifluoroacetic acid (TFA) and formic acid were added to the digested peptides for a final concentration of 0.2% (pH ∼2). Peptides were desalted with 50-mg Sep-Pak C18 columns (Waters). De-salted protein was resuspended in 100 mM triethylammonium bicarbonate (TEAB) and labeled with mass tags using the Thermo TMT10plex protocol. After labeling and quenching, each labeled IP condition was multiplexed together and de-salted again to remove excess TMT label. The final peptide mixture was resuspending in 0.1% TFA in preparation for LC-MS/MS analysis using a Fusion Lumos mass spectrometer.

Relative changes in peptide concentration were determined at the MS3 level by isolating and fragmenting the five most dominant MS2 ion peaks. Raw files were searched using the Andromeda engine embedded in MaxQuant (v.2). Variable modifications included oxidation (M) and protein amino-terminal acetylation. Carbamidomethyl (C) was a fixed modification. The number of modifications per peptide was capped at five. Digestion was set to tryptic (proline blocked). The 1% false recovery rate (FDR) was determined using a reverse decoy proteome.

### In silico protein structure prediction

Structure of the protein complex containing human Cyclin D1, CDK6 and Rb helix (amino acids 895-915) was predicted using AlphaFold2 (v. 1.5.2). Multiple sequence alignments were generated using the ColabFold pipeline, and structures were predicted with a maximum of 30 recycles and 4 predictions per model, generating a total of 20 predictions. The top-ranked model was subjected to Amber relaxation and was selected for further analysis and visualization using UCSF ChimeraX (v. 1.7.1). Contacts between chains were determined using the Contacts tool, with van der Waals overlap of at least 0.40 A.

### Peptide binding assays using SIMBA

Cyclin-peptide binding was measured by Systematic Intracellular Motif Binding Analysis (SIMBA), a yeast-based binding assay^40,57^. For low-throughput tests, galactose-inducible cyclin D1-Cln2 fusion constructs were co-transformed into yeast strain PPY2617 with plasmids encoding Ste20^Ste5PM^ chimeras that harbored a control peptide (DVCTPR), an RxL peptide (VSAVRKLDFSDT), or an RBL1 peptide (DVLLKRLQDVVSERAN). Binding of the cyclin D1 fusion to the peptide sequence inhibits pheromone signaling and activation of a transcription reporter (*P_FUS1_-lacZ*). Cell cultures were grown in synthetic complete (SC) liquid medium with 2% raffinose lacking histidine and uracil. Expression of the cyclin D1-Cln2 fusion was induced by adding 2% galactose for 90 minutes, then cells were treated with *α*-factor (50 nM) for 45 min, and then ꞵ-galactosidase activity was measured in cell lysates.

High-throughput SIMBA experiments used competitive growth methods described previously^40,57^. Plasmid libraries were constructed to encode all single-residue variants of 16-residue RBL1 and RBL2 peptide sequences, inserted into a Ste20^Ste5PM^ chimera construct. These sequences were amplified by PCR using oligonucleotides as templates and primers (actatacgcgtGGAGGTAGTGGT and tctttgcatgcTCCAGATCCACC) that anneal to flanking linker sequences (Gly Gly Ser Gly). In the template oligonucleotides, single codons were randomized (NNN) to generate all 64 nucleotide variants. PCR products were digested with restriction enzymes (MluI and SphI), treated with calf intestinal phosphatase, and ligated with a digested Ste20^Ste5PM^ vector (pPP4745). Ligation products were transformed into E. coli (XL-10 Gold Ultracompetent Cells; Agilent/Agilent Technologies) and plated on LB+Amp plates, and then plasmid DNA was purified from suspended colonies. Separately, 13-residue variants of RBL1, RBL2, and Rb were included in a previously-described library (RxL library 2) that was used to analyze cyclin binding to RxL motif peptides^57^. The plasmid libraries were transformed into yeast strains containing integrated, galactose-inducible constructs expressing yeast Cln2 fused to cyclins D1, D2, or D3^57^. Transformation mixtures were plated on SC–URA medium and incubated for 3 days. Colonies were suspended in –URA/raffinose liquid medium, washed, diluted into fresh medium, incubated in a shaking water bath for ∼24 hrs, then treated with 2% galactose (75 min) to induce cyclin expression. At this time (t = 0), cell aliquots were harvested, and the remaining cultures were treated with *α*-factor (500 nM) and returned to incubation, with additional aliquots harvested at 8, 20, and 32 hr. Harvested cells were collected by centrifugation, washed and resuspended with water, then transferred to microcentrifuge tubes. These suspensions were centrifuged, the supernatants were aspirated, and the pellets were frozen using liquid nitrogen and stored at −80°C. Plasmid DNAs were isolated from frozen yeast cell pellets as described previously^40,57^ using a Zymo Research ZR Plasmid Miniprep Kit (#D4015). Samples of recovered plasmid DNA were amplified by PCR using primers that include standard P5 and P7 sequences for binding to Illumina flow cells during next generation sequencing (NGS). Aliquots of the PCR products were analyzed by agarose gel electrophoresis, band intensities were quantified by densitometry (ImageJ), and then equal amounts of products from all timepoints for a given strain were pooled, purified using Zymo Spin I columns (Zymo Research #C1003-250), and eluted in 30 μL of 10 mM TRIS-HCl, pH 8. The concentration of the eluted products was measured using NanoDrop spectrophotometry, and then equal amounts of samples from all strains were pooled and gel-purified. The final sample was sent for Illumina-based NGS (paired-end sequencing, 150 bp) by a commercial vendor (Novogene).

NGS data were de-multiplexed using bar code and index identifiers for each strain and timepoint. To analyze mutational scanning of RBL1 and RBL2 sequences, we used Enrich2 software^65^ to obtain read counts for all sequence variants, and added 0.5 counts to any variants depleted to 0 counts by the 32-hr timepoint. Then, we used Enrich2 to calculate fitness scores that describe the rate of change of mutant variants compared to the wild-type sequence. Separately, variants of RBL1, RBL2, and Rb sequences in the earlier-reported library (RxL library 2) were analyzed as described previously^57^. To create logos of sequence preferences, we transformed Enrich2 scores into a preference metric that expresses the bias for each residue at a given motif position relative to the set of all possible residues, as described previously^40,57^. Calculation of Gini coefficients from mutational scanning data began with normalizing the raw Enrich2 score for each amino acid variant to the lowest (= 0) and highest (= 1) scores in a given motif array. Gini coefficients were calculated in Microsoft Excel for each position of a parent peptide, separately for each of 6 pair-wise combinations of 3 cyclins (D1, D2, D3) and 2 sets of peptide mutations (RBL1, RBL2). Then, the average of all 6 values at each peptide position was plotted. To compare Gini coefficients to predicted interface residues, AlphaFold3^66^ was used to predict structures of complexes between 3 cyclins (D1, D2, D3) and each of 3 peptides (Rb, RBL1, RBL2). The models of cyclin D2 with Rb and RBL2 were discarded due to placement of the peptide into the hydrophobic patch. For the remaining 7 complexes, the solvent-accessible surface areas of peptide residues in the bound and unbound states were extracted using Chimera X software^67^, and the buried fraction was calculated.

### Cyclin structural alignment

Structural alignment of cyclin structures was performed in UCSF ChimeraX using the Matchmaker tool. Structures from PDB were aligned using default Matchmaker settings with the cyclin D1 chain of 2W96 as the reference and pairwise sequence alignment enabled. The resulting structure-guided sequence alignment was analyzed and exported from the ChimeraX Sequence Viewer.

### Protein expression and purification

Full-length, N-terminally glutathione S-transferase (GST)–tagged retinoblastoma protein (Rb) was expressed in *Escherichia coli* and purified by glutathione–agarose affinity chromatography as described previously^24^. Briefly, the pGEX-4T1-GST-Rb plasmid was transformed into *E. coli* BL21(DE3)RIL cells (Agilent, #230245), and protein expression was induced with 0.1 mM IPTG overnight at 18 °C. Cells were harvested and lysed in lysis buffer (50 mM Tris, pH 8.0; 100 mM NaCl; 0.5 mM EDTA; 0.5 mM EGTA) supplemented with 1 mg/mL lysozyme, 50 µg/mL DNase, 1 mM PMSF, 1 µg/mL pepstatin A, 1 µg/mL aprotinin, and 1 µg/mL leupeptin. Lysates were incubated at 4 °C for 10 min with stirring and subsequently disrupted by sonication (2 × 15 s). Immediately after sonication, the NaCl concentration was increased to 500 mM, and insoluble material was removed by centrifugation. The cleared lysate supplemented with 0.1% Tween-20 was incubated with 200 µL glutathione–agarose beads (Sigma-Aldrich, G4510) for 2 h at 4 °C with gentle mixing. Beads were loaded onto columns (BioRad, 731-1550) and washed with 10 column volumes of wash buffer (50 mM Tris, pH 8.0; 100 mM potassium acetate; 25 mM magnesium acetate; 0.1% Tween-20), followed by 5 column volumes of wash buffer containing 1 mM ATP. GST-Rb was eluted in four fractions using an elution buffer (50 mM Tris, pH 8.0; 100 mM potassium acetate; 25 mM magnesium acetate; 10% glycerol; 15 mM reduced glutathione). Protein purity and yield were assessed by Coomassie staining.

Non-docking, Helix-docking, and RxL-docking GST–Cdk reporter substrate proteins, each containing an N-terminal GST tag, a TEV protease cleavage site and a peptide derived from Rb containing a single Cdk phosphorylation site (Kitagawa et al., 1996), were expressed and purified as described above. For the Helix-docking and RxL-docking reporters, docking sequences corresponding to the Rb C-terminal helix (Rb residues 895–915: SKFQQKLAEMTSTRTRMQKQK) or the Cdc6 RxL docking motif (Cdc6 residues 89–103: HTLKGRRLVFDNQLT) ^68^ were fused to the C terminus of the proteins via a G4S glycine–serine linker (GGGGS).

An N-terminally 6×His-tagged C-terminal fragment of Rb (RbC; amino acids 772–928) was expressed in E. coli BL21(DE3)RIL (Agilent, #230245) cells and purified by cobalt affinity chromatography. Proteins were eluted in a buffer containing 25 mM HEPES (pH 7.4), 300 mM NaCl, 10% glycerol, and 200 mM imidazole.

Human cyclin–Cdk fusion complexes were purified from Saccharomyces cerevisiae as previously described^14,24,69^. Briefly, pRS425-pGAL1-3×FLAG–cyclin D–L–Cdk4 or Cdk6 plasmids (where L denotes a 3×GGGGS linker) were transformed into yeast, and protein expression was induced at an OD₆₀₀ of 0.6–0.8, followed by incubation at 30 °C for 3 h. Cells were harvested, and cyclin–Cdk complexes were purified using anti-FLAG M2 affinity agarose beads (Sigma-Aldrich, #A2220) and eluted with elution buffer containing 50 mM HEPES-KOH (pH 7.6), 250 mM KCl, 1 mM MgCl₂, 1 mM EGTA, 5% glycerol, and 0.2 mg/mL 3×FLAG peptide (Sigma-Aldrich, #F4799). Purified cyclin–Cdk complexes were verified by Western blotting using an anti-FLAG antibody (1:2,000; Sigma-Aldrich, #F1804) and a donkey anti-mouse IRDye 800CW secondary antibody (1:10,000; LI-COR, #926-32212).

Cyclin D1–Cdk4–p27(pY) trimer complexes used in this study were a gift from the Rubin laboratory^7^.

### *In vitro* kinase assays

For all experiments, substrate concentrations ranged from 0.5–5 µM across different experiments but were held constant within each experiment. Enzyme concentrations were maintained in the low nanomolar range and were chosen such that less than 10% of the substrate was converted by the end of the experiment, unless stated otherwise. Reaction aliquots were collected at two time points (typically 8 and 16 min, unless stated otherwise), and reactions were terminated by the addition of SDS–PAGE sample buffer.

*In vitro* kinase assays using radiolabeled [γ-³²P]ATP (Fig 1C-F, 4B) were performed in a basal reaction mixture containing 20 mM Tris (pH 8.0), 150 mM NaCl, 5 mM MgCl₂, 10 mM magnesium acetate, 40 mM potassium acetate, 6 mM glutathione, 0.2 mg/mL 3×FLAG peptide, 6% glycerol, 3 mM EGTA, 0.2 mg/mL BSA, and 500 µM ATP, supplemented with 2 µCi [γ-³²P]ATP per reaction (PerkinElmer, BLU502Z250UC). Phosphorylated proteins were resolved on 6-10% SDS–PAGE gels and visualized by autoradiography using a Typhoon 9210 imager (GE Healthcare Life Sciences). Autoradiographs were quantified using ImageQuant TL software (v10.2-499).

*In vitro* kinase assays using ATP-γ-S (Fig. 4C and D) were performed as previously described^54^, with minor modifications. Reactions contained 50 mM HEPES (pH 7.4), 150 mM NaCl, 5 mM MgCl₂, 0.5 mM DTT, 0.08 mg/mL BSA, 0.5 mM ATP-γ-S, and FLAG elution buffer to normalize reaction volumes across different cyclin–Cdk complexes. Throughout the kinase reactions, His-RbC (Rb aa: 772–928) was used at low micromolar concentrations (0.5–2 µM), while enzyme concentrations were maintained in the low nanomolar range. Kinase reactions were incubated at room temperature for 8 or 16 min and then quenched by the addition of EDTA to a final concentration of 20 mM. Thiophosphorylated substrates were subsequently alkylated with 2.5 mM p-nitrobenzyl mesylate (pNBM; Selleckchem, #E1248) for 1 h at room temperature. Alkylation reactions were terminated by the addition of 20 mM DTT and 1× Laemmli sample buffer (Bio-Rad, #1610747), followed by heating at 70 °C for 10 min and centrifugation at 16,000 × g for 1 min. Samples were analyzed by SDS–PAGE followed by Western blotting. Total protein was visualized using the Revert™ 700 Total Protein Stain Kit (LI-COR, #926-11010). Primary antibodies used were rabbit anti–thiophosphate ester (1:2,000; Abcam, #ab133473) and mouse anti-FLAG (1:5,000; Sigma-Aldrich, #F1804). Secondary antibodies were donkey anti-rabbit IRDye 800CW and donkey anti-mouse IRDye 680CW (1:10,000; LI-COR, #926-32213 and #926-68022, respectively). Fluorescent signals were detected using an Amersham Typhoon imager (Cytiva) and quantified with ImageQuant TL software (v10.2-499).

### Peptide synthesis

Peptides were synthesized by Pepmic Co., Ltd and Anaspec. Lyophilized peptides were resuspended in 50 mM HEPES pH 7.4. Rb Helix peptide sequence is SKFQQKLAEMTSTRTRMQKQK. The RxL peptide sequence is RVNRILFPT.

## Acknowledgements

We thank all the members of the Koivomagi and Skotheim labs for helpful comments on the manuscript. This work was primarily supported by the NIH NCI through P01 CA254867 to JMS. This work utilized the computational resources of the NIH HPC Biowulf cluster (https://hpc.nih.gov). This research was also supported, in part, by the Intramural Research Program of the National Institutes of Health (NIH Grant ZIA BC 012133 to M.K.) and a grant from the NIH to P.M.P. (R01GM145795). M.O. and N.E.D. were funded by a Cancer Research UK Senior Cancer Research Fellowship (C68484/A28159) and M.O. was also funded by a UKRI grant EP/X042065/1. The contributions of the NIH author(s) are considered Works of the United States Government. The findings and conclusions presented in this paper are those of the author(s) and do not necessarily reflect the views of the NIH or the U.S. Department of Health and Human Services.

## References

1. Fassl, A., Geng, Y. & Sicinski, P. CDK4 and CDK6 kinases: From basic science to cancer therapy. Science 375, eabc1495 (2022).

2. Pennycook, B. R. & Barr, A. R. Restriction point regulation at the crossroads between quiescence and cell proliferation. FEBS Lett 594, 2046–2060 (2020).

3. Rubin, S. M., Sage, J. & Skotheim, J. M. Integrating Old and New Paradigms of G1/S Control. Mol Cell 80, 183–192 (2020).

4. Malumbres, M. & Barbacid, M. Cell cycle, CDKs and cancer: a changing paradigm. Nat Rev Cancer 9, 153–166 (2009).

5. Morgan, D. O. The Cell Cycle: Principles of Control. (Published by New Science Press in association with Oxford University Press; Distributed inside North America by Sinauer Associates, Publishers, London: Sunderland, MA, 2007).

6. Sherr, C. J. & Roberts, J. M. CDK inhibitors: positive and negative regulators of G1-phase progression. Genes Dev 13, 1501–1512 (1999).

7. Guiley, K. Z. et al. p27 allosterically activates cyclin-dependent kinase 4 and antagonizes palbociclib inhibition. Science 366, eaaw2106 (2019).

8. Bertoli, C., Skotheim, J. M. & de Bruin, R. A. M. Control of cell cycle transcription during G1 and S phases. Nat Rev Mol Cell Biol 14, 518–528 (2013).

9. Kent, L. N. & Leone, G. The broken cycle: E2F dysfunction in cancer. Nat Rev Cancer 19, 326–338 (2019).

10. Trimarchi, J. M. & Lees, J. A. Sibling rivalry in the E2F family. Nat Rev Mol Cell Biol 3, 11–20 (2002).

11. Dick, F. A. & Rubin, S. M. Molecular mechanisms underlying RB protein function. Nat Rev Mol Cell Biol 14, 297–306 (2013).

12. Merrick, K. A. et al. Switching Cdk2 on or off with small molecules to reveal requirements in human cell proliferation. Mol Cell 42, 624–636 (2011).

13. Pardee, A. B. A restriction point for control of normal animal cell proliferation. Proc Natl Acad Sci U S A 71, 1286–1290 (1974).

14. Schwarz, C. et al. A Precise Cdk Activity Threshold Determines Passage through the Restriction Point. Mol Cell 69, 253–264.e5 (2018).

15. Narasimha, A. M. et al. Cyclin D activates the Rb tumor suppressor by mono-phosphorylation. Elife 3, (2014).

16. Sanidas, I. et al. A Code of Mono-phosphorylation Modulates the Function of RB. Mol Cell 73, 985–1000.e6 (2019).

17. Musgrove, E. A., Caldon, C. E., Barraclough, J., Stone, A. & Sutherland, R. L. Cyclin D as a therapeutic target in cancer. Nature Reviews Cancer 11, 558–572 (2011).

18. Zatulovskiy, E., Zhang, S., Berenson, D. F., Topacio, B. R. & Skotheim, J. M. Cell growth dilutes the cell cycle inhibitor Rb to trigger cell division. Science 369, 466–471 (2020).

19. Zhang, S. et al. The G1-S transition is promoted by Rb degradation via the E3 ligase UBR5. Science Advances 10, eadq6858 (2024).

20. Chung, M. et al. Transient Hysteresis in CDK4/6 Activity Underlies Passage of the Restriction Point in G1. Mol Cell 76, 562–573.e4 (2019).

21. Yang, H. W. et al. Stress-mediated exit to quiescence restricted by increasing persistence in CDK4/6 activation. Elife 9, e44571 (2020).

22. Minireview: Cyclin D1: Normal and Abnormal Functions | Endocrinology | Oxford Academic. https://academic.oup.com/endo/article-abstract/145/12/5439/2499715?redirectedFrom=fulltext&login=false.

23. Dowdy, S. F. et al. Physical interaction of the retinoblastoma protein with human D cyclins. Cell 73, 499–511 (1993).

24. Topacio, B. R. et al. Cyclin D-Cdk4,6 Drives Cell-Cycle Progression via the Retinoblastoma Protein’s C-Terminal Helix. Mol Cell 74, 758–770.e4 (2019).

25. Petre, C. E., Wetherill, Y. B., Danielsen, M. & Knudsen, K. E. Cyclin D1: Mechanism and Consequence of Androgen Receptor Co-repressor Activity *. Journal of Biological Chemistry 277, 2207–2215 (2002).

26. Zwijsen, R. M. et al. CDK-independent activation of estrogen receptor by cyclin D1. Cell 88, 405–415 (1997).

27. Zwijsen, R. M., Buckle, R. S., Hijmans, E. M., Loomans, C. J. & Bernards, R. Ligand-independent recruitment of steroid receptor coactivators to estrogen receptor by cyclin D1. Genes Dev 12, 3488–3498 (1998).

28. Schulman, B. A., Lindstrom, D. L. & Harlow, E. Substrate recruitment to cyclin-dependent kinase 2 by a multipurpose docking site on cyclin A. Proc Natl Acad Sci U S A 95, 10453–10458 (1998).

29. Wood, D. J. & Endicott, J. A. Structural insights into the functional diversity of the CDK-cyclin family. Open Biol 8, 180112 (2018).

30. Blain, S. W., Montalvo, E. & Massagué, J. Differential interaction of the cyclin-dependent kinase (Cdk) inhibitor p27Kip1 with cyclin A-Cdk2 and cyclin D2-Cdk4. J Biol Chem 272, 25863–25872 (1997).

31. Cheng, M. et al. The p21(Cip1) and p27(Kip1) CDK ‘inhibitors’ are essential activators of cyclin D-dependent kinases in murine fibroblasts. EMBO J 18, 1571–1583 (1999).

32. James, M. K., Ray, A., Leznova, D. & Blain, S. W. Differential modification of p27Kip1 controls its cyclin D-cdk4 inhibitory activity. Mol Cell Biol 28, 498–510 (2008).

33. LaBaer, J. et al. New functional activities for the p21 family of CDK inhibitors. Genes Dev 11, 847–862 (1997).

34. Adams, P. D. et al. Retinoblastoma protein contains a C-terminal motif that targets it for phosphorylation by cyclin-cdk complexes. Mol Cell Biol 19, 1068–1080 (1999).

35. Hirschi, A. et al. An overlapping kinase and phosphatase docking site regulates activity of the retinoblastoma protein. Nat Struct Mol Biol 17, 1051–1057 (2010).

36. Bhaduri, S. et al. A Docking Interface in the Cyclin Cln2 Promotes Multi-site Phosphorylation of Substrates and Timely Cell-Cycle Entry. Current Biology 25, 316–325 (2015).

37. Loog, M. & Morgan, D. O. Cyclin specificity in the phosphorylation of cyclin-dependent kinase substrates. Nature 434, 104–108 (2005).

38. Örd, M., Venta, R., Möll, K., Valk, E. & Loog, M. Cyclin-Specific Docking Mechanisms Reveal the Complexity of M-CDK Function in the Cell Cycle. Mol Cell 75, 76–89.e3 (2019).

39. Örd, M. et al. Proline-Rich Motifs Control G2-CDK Target Phosphorylation and Priming an Anchoring Protein for Polo Kinase Localization. Cell Rep 31, 107757 (2020).

40. Subbanna, M. S., Winters, M. J., Örd, M., Davey, N. E. & Pryciak, P. M. A quantitative intracellular peptide-binding assay reveals recognition determinants and context dependence of short linear motifs. J Biol Chem 301, 108225 (2025).

41. Rubin, S. M., Gall, A.-L., Zheng, N. & Pavletich, N. P. Structure of the Rb C-terminal domain bound to E2F1-DP1: a mechanism for phosphorylation-induced E2F release. Cell 123, 1093–1106 (2005).

42. S, B., et al. The soft agar colony formation assay. PubMed https://pubmed.ncbi.nlm.nih.gov/25408172/ (2014).

43. Bienvenu, F. et al. Transcriptional role of cyclin D1 in development revealed by a genetic– proteomic screen. Nature 463, 374–378 (2010).

44. Schulman, B. A., Lindstrom, D. L. & Harlow, E. Substrate recruitment to cyclin-dependent kinase 2 by a multipurpose docking site on cyclin A. Proceedings of the National Academy of Sciences 95, 10453–10458 (1998).

45. M, M. & E, H. Identification of G1 kinase activity for cdk6, a novel cyclin D partner. PubMed https://pubmed.ncbi.nlm.nih.gov/8114739/.

46. Afonso, A. L., Cavaleiro, C. T., Castanho, M. A. R. B., Neves, V. & Cavaco, M. The Potential of Peptide-Based Inhibitors in Disrupting Protein–Protein Interactions for Targeted Cancer Therapy. International Journal of Molecular Sciences 26, 3117 (2025).

47. Örd, M. et al. Uncovering cancer dependencies in peptide-interacting protein pockets. 2026.03.13.711608 Preprint at 10.64898/2026.03.13.711608 (2026).

48. Zhao, H., Shen, A., Xiang, Y. K. & Corey, D. P. Three Recombinant Engineered Antibodies against Recombinant Tags with High Affinity and Specificity. PLOS ONE 11, e0150125 (2016).

49. Örd, M. et al. Multisite phosphorylation code of CDK. Nat Struct Mol Biol 26, 649–658 (2019).

50. Stern, B. & Nurse, P. A quantitative model for the cdc2 control of S phase and mitosis in fission yeast. Trends in Genetics 12, 345–350 (1996).

51. Swaffer, M. P., Jones, A. W., Flynn, H. R., Snijders, A. P. & Nurse, P. CDK Substrate Phosphorylation and Ordering the Cell Cycle. Cell 167, 1750–1761.e16 (2016).

52. Zhang, L. et al. Multiple Layers of Phospho-Regulation Coordinate Metabolism and the Cell Cycle in Budding Yeast. Front. Cell Dev. Biol. 7, 338 (2019).

53. Kõivomägi, M. et al. Dynamics of Cdk1 Substrate Specificity during the Cell Cycle. Molecular Cell 42, 610–623 (2011).

54. W, Z., et al. Distinct Allosteric Networks in CDK4 and CDK6 in the Cell Cycle and in Drug Resistance. PubMed https://pubmed.ncbi.nlm.nih.gov/40174666/ (2026).

55. Kõivomägi, M., Swaffer, M. P., Turner, J. J., Marinov, G. & Skotheim, J. M. G1 cyclin-Cdk promotes cell cycle entry through localized phosphorylation of RNA polymerase II. Science 374, 347–351 (2021).

56. Archambault, V., Buchler, N. E., Wilmes, G. M., Jacobson, M. D. & Cross, F. R. Two-faced cyclins with eyes on the targets. Cell Cycle 4, 125–130 (2005).

57. Örd, M. et al. High-throughput investigation of cyclin docking interactions reveals the complexity of motif binding determinants. Nature Communications 16, 7622 (2025).

58. Hafner, M. et al. Multiomics Profiling Establishes the Polypharmacology of FDA-Approved CDK4/6 Inhibitors and the Potential for Differential Clinical Activity. Cell Chemical Biology 26, 1067–1080.e8 (2019).

59. Sherr, C. J., Beach, D. & Shapiro, G. I. Targeting CDK4 and CDK6: From Discovery to Therapy. Cancer Discov 6, 353–367 (2016).

60. Slamon, D. J. et al. Overall Survival with Ribociclib plus Fulvestrant in Advanced Breast Cancer. N Engl J Med 382, 514–524 (2020).

61. Turner, N. C. et al. Overall Survival with Palbociclib and Fulvestrant in Advanced Breast Cancer. N Engl J Med 379, 1926–1936 (2018).

62. Rubin, S. M., Sage, J. & Skotheim, J. M. Emerging Strategies to Inhibit the G1/S Transition for Cancer Therapy. Cancer Res 10.1158/0008-5472.CAN-25-0916 (2026) doi:10.1158/0008-5472.CAN-25-0916.

63. Kl, M., et al. The pINDUCER lentiviral toolkit for inducible RNA interference in vitro and in vivo. PubMed https://pubmed.ncbi.nlm.nih.gov/21307310/ (2011).

64. Fa, D., E, S. & Nj, D. Mutagenesis of the pRB pocket reveals that cell cycle arrest functions are separable from binding to viral oncoproteins. PubMed https://pubmed.ncbi.nlm.nih.gov/10779361/.

65. Rubin, A. F. et al. A statistical framework for analyzing deep mutational scanning data. Genome Biol 18, 150 (2017).

66. Abramson, J. et al. Accurate structure prediction of biomolecular interactions with AlphaFold 3. Nature 630, 493–500 (2024).

67. Meng, E. C. et al. UCSF ChimeraX: Tools for structure building and analysis. Protein Sci 32, e4792 (2023).

68. Takeda, D. Y., Wohlschlegel, J. A. & Dutta, A. A Bipartite Substrate Recognition Motif for Cyclin-dependent Kinases. Journal of Biological Chemistry 276, 1993–1997 (2001).

69. M, K. Purification of Cyclin-Dependent Kinase Fusion Complexes for In Vitro Analysis. PubMed https://pubmed.ncbi.nlm.nih.gov/34085218/.

