## Supplemental Figures for "Identification and inhibition of the Cyclin D Rb-docking interface that drives cell division"

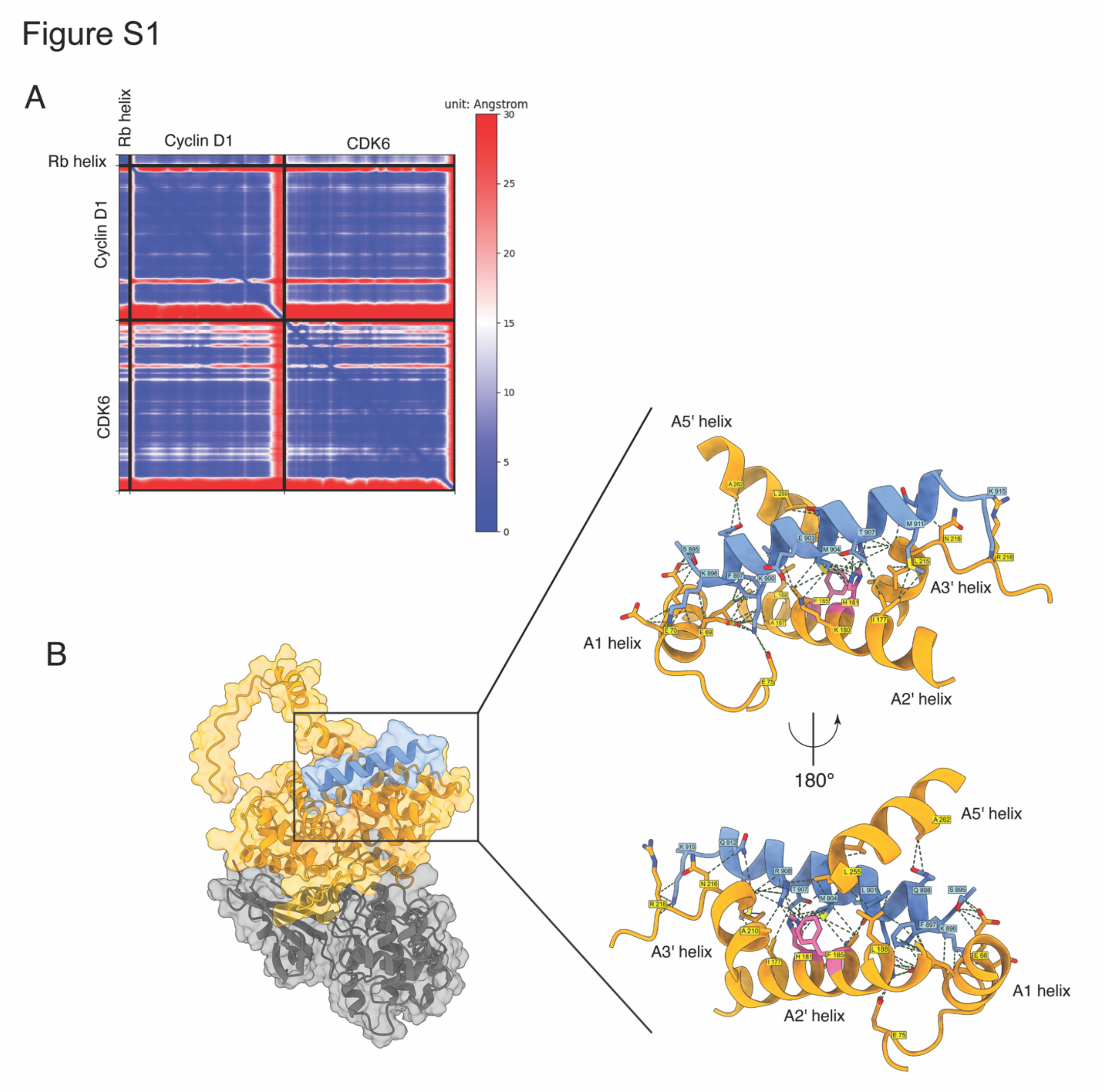


**Figure S1: Cyclin D's A2' helix docks Rb to target it for phosphorylation.**

A. Prediction aligned error (PAE) scores for the top-ranked Rb helix-cyclin D1-CDK6 multimeric model. Axes indicate the position of individual amino acids for the denoted protein.

B. Model showing the Rb helix (blue), cyclin D1 (yellow) and CDK6 (grey) complex. Inset shows the interaction interface between the Rb helix and Cyclin D1. H181 and F185 are denoted as pink as the main cyclin D1 interface residues mutated in the present study.

Figure S2

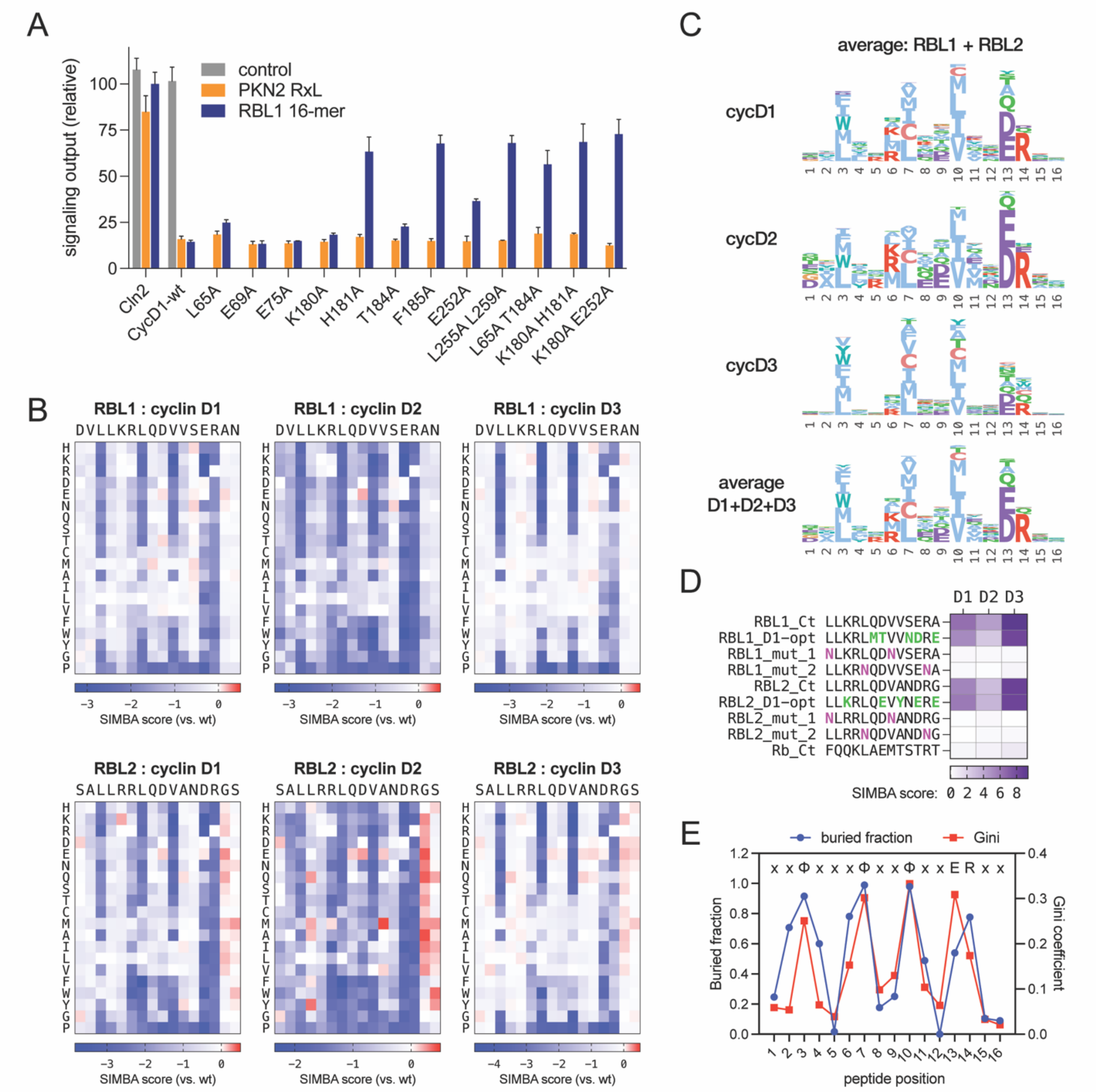

**Figure S2: Deep mutational scanning of cyclin D docking helices reveals the binding determinants.**

A. Binding of different cyclin D1 mutants to RBL1 helix and an RxL motif from PKN2 measured in SIMBA using a pFUS1-lacZ reporter as a readout for docking-dependent suppression of yeast mating signaling. Bars, mean ± standard deviation (SD); n = 4 biological replicates.

B. Heatmaps showing the wild-type normalized binding strengths of all single mutation variants of RBL1 and RBL2 peptides measured in pooled SIMBA experiments.

C. Sequence logos showing the favored residues at each position for cyclin D1, D2, D3 and the averaged results of all three D-type cyclins.

D. SIMBA scores indicating binding strengths of wild-type and mutant RBL1, RBL2 and Rb helices to cyclin D1, D2, and D3.
E. Residue selectivity in the SIMBA DMS data was summarized by calculating the Gini coefficient; six combinations (3 cyclins x 2 peptides) were averaged. The fraction of buried surface area was calculated from AlphaFold3 models of cyclins D1, D2, and D3 bound to RBL1, RBL2, and Rb helices; the average is plotted.

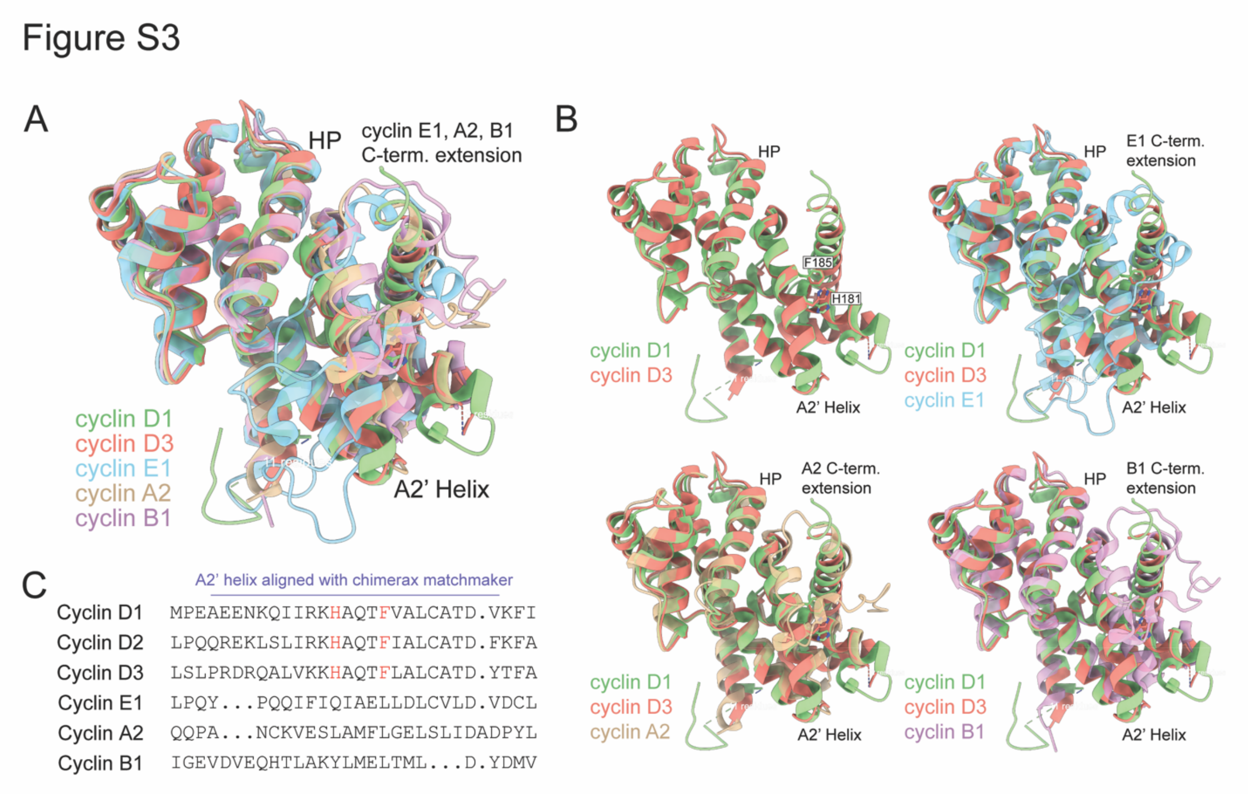


**Figure S3: Cyclin D's A2' helix docks Rb to target it for phosphorylation.**

A. Alignment of cyclin D3 (red, PDB: 3G33), E1 (cyan, PDB: 1W98), A2 (brown, PDB: 1JST), and B1 (magenta, PDB: 2JGZ) against cyclin D1 (green, PDB: 2W96) as the reference sequence.
B. Alignment of D-type cyclins alone, or against the individual cyclins E1, A2, or B1.
C. A sequence comparison of the residues comprising the A2' helix of the D-type cyclins with the other cell cycle cyclins E1, A2, and B1 based on structural alignment with Matchmaker tool in ChimeraX. Cyclins E1, A2, and B1 do not have the conserved interface residues of the cyclin D1 A2' helix (H181 and F185).

Figure S4

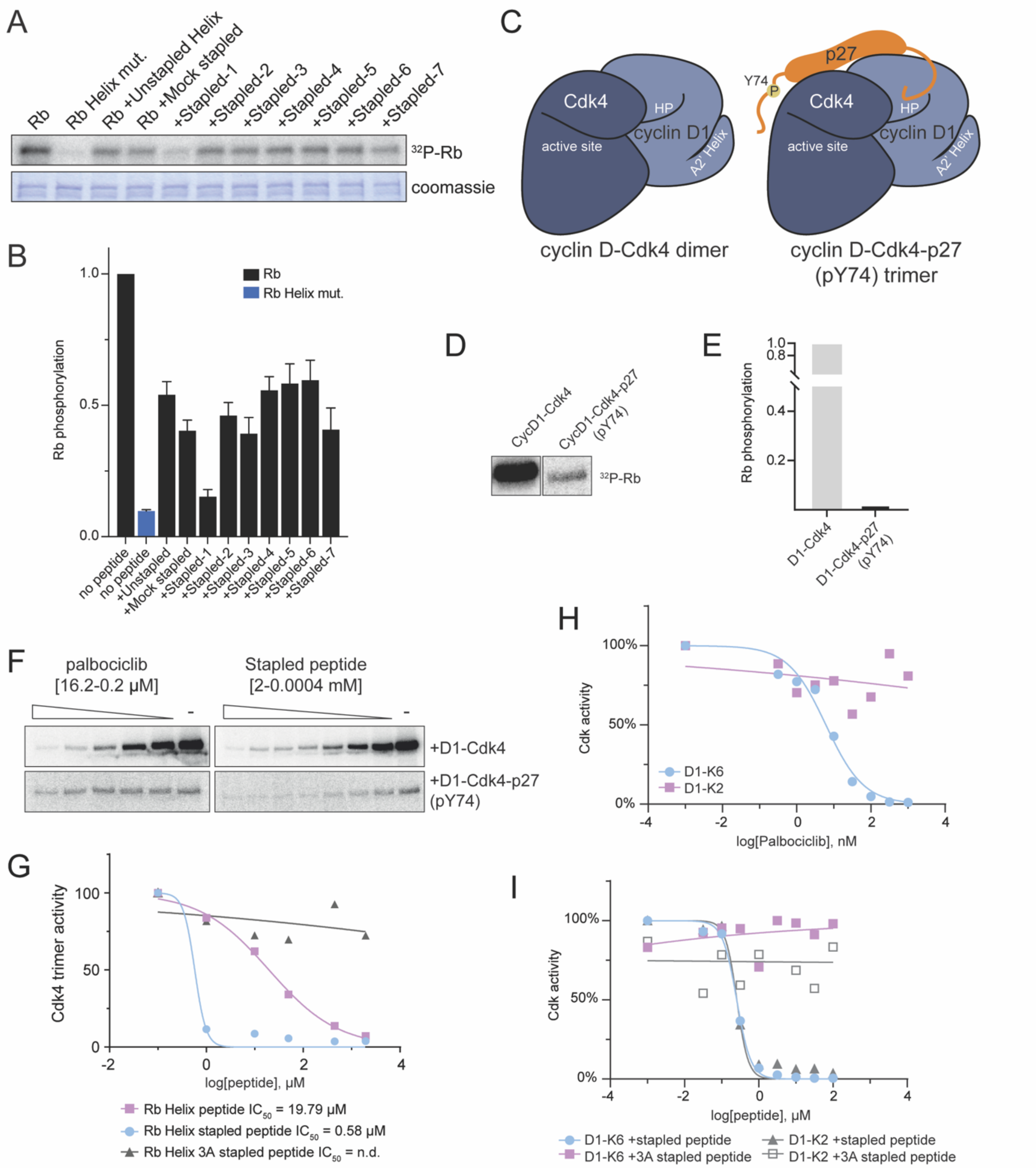


**Figure S4: Rb C-terminal peptides can inhibit the cyclin D A2' Helix in dimer, trimer, and non-canonical complexes.**

A. Representative *in vitro* kinase assays with full length Rb variant and cyclin D1-CDK4 in the presence of (concentration) the indicated stapled or unstapled Rb Helix peptide. RbWT denotes wild-type Rb and RbHelix mut. denotes an Rb variant where the helix-based docking interface residues F897, L901, and R908 are substituted with alanines.

B. Quantification of *in vitro* kinase assays as in A (n=3).
C. Schematic showing the cyclin D-CDK4 dimer and cyclin D-CDK4-p27 trimer complexes.
D. Representative *in vitro* kinase assay with full length Rb comparing cyclin D1-CDK4 dimer and cyclin D1-CDK4-p27 trimer complexes.
E. Quantification of *in vitro* kinase assays as in B (n=3).
F. Representative *in vitro* kinase assays using cyclin D1-CDK4-p27 trimer complex and full length Rb protein. Rb-derived stapled helix peptide was added to the kinase reactions at the indicated concentrations.
G. Quantification of *in vitro* kinase assays using cyclin D1-CDK4-p27 trimer complex and full length Rb protein as in F (n=3).
H. Quantification of *in vitro* kinase assays using cyclin D1-CDK6 or cyclin D1-CDK2 and full length Rb proteins with palbociclib.
I. Quantification of *in vitro* kinase assays using cyclin D1-CDK6 or cyclin D1-CDK2 and full length Rb protein. Rb-derived stapled helix peptides or a similar stapled peptide with the cyclin D-docking residues substituted with alanines (3A) were added to the kinase reactions at the indicated concentrations.

Figure S5

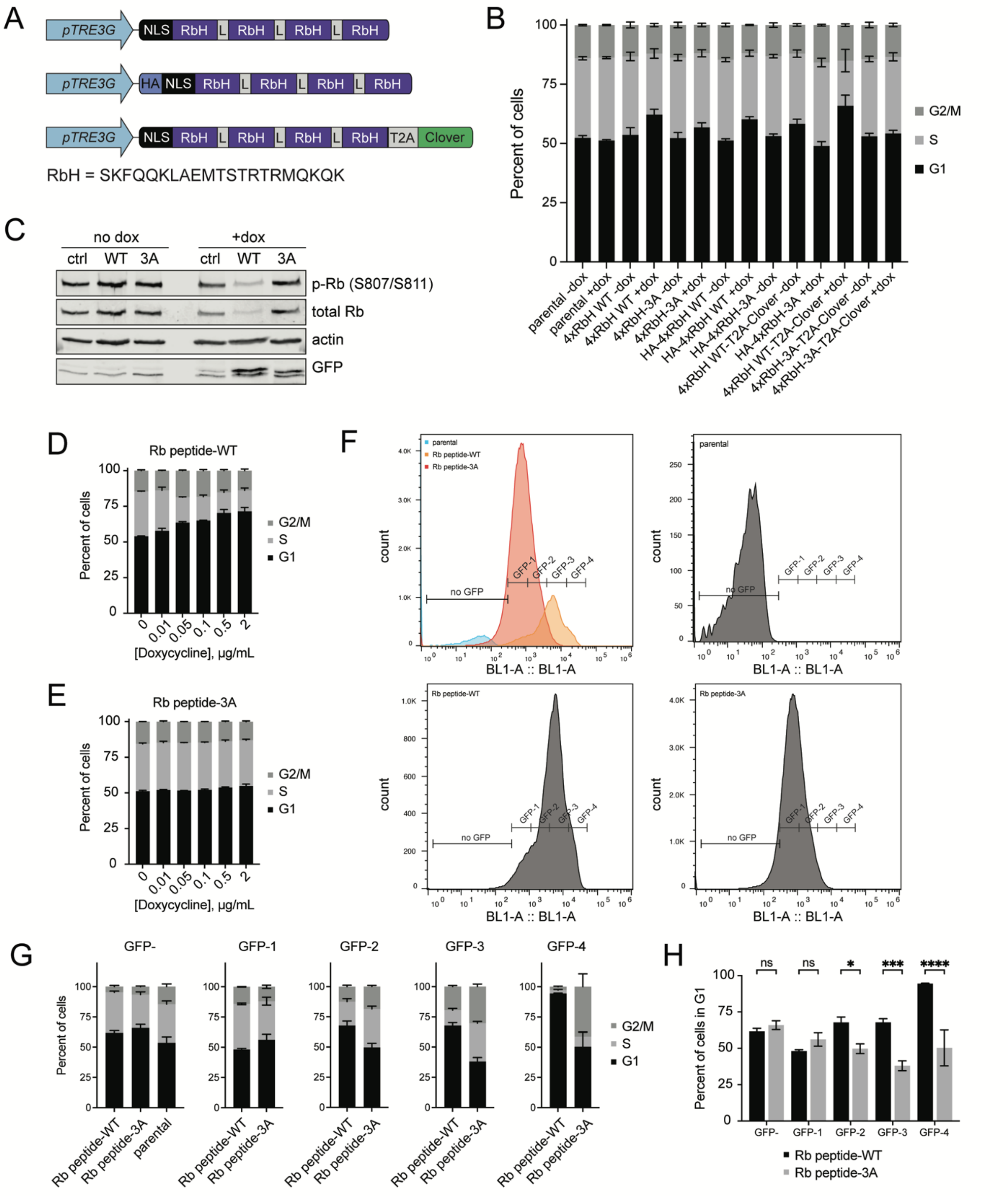


**Figure S5: The wildtype Rb C-terminal helix peptide inhibits proliferation in cells.**

A. PiggyBac integration constructs containing doxycycline-inducible NLS-4xRbHelix, HA-NLS-4xRbHelix, or NLS-4xRbHelix-T2A-Clover. Constructs were integrated into RPE-1 cells. Rb Helix sequence is shown.

B. Flow cytometry analysis of cell cycle phase distribution using EdU incorporation and DAPI staining of control RPE-1 cells (parental) or RPE-1 cells expressing WT or 3A variants of the 4xRbHelix constructs described in Fig. S5A after 48 hours of doxycycline treatment. 3A denotes a 4xRbHelix variant in which the 3 interface residues were substituted with alanines within all four helix sequences. n = 3.
C. Representative immunoblot analysis of lysates from cells containing the NLS-4xRbHelix-Clover construct indicated in Fig. 5C after 48 hours of doxycycline treatment.
D. Flow cytometry analysis of cell cycle phase distribution using EdU incorporation and DAPI staining of cells expressing Rb peptide-WT after 48 hours of 0 to 2 μg/mL doxycycline. n= 3.
E. Flow cytometry analysis of cell cycle phase distribution using EdU incorporation and DAPI staining of cells expressing Rb peptide-3A after 48 hours of 0 to 2 μg/mL doxycycline. n= 3.
F. Representative combined or individual histograms showing GFP-negative and four bins of GFP-positive expression in parental or Rb peptide-expressing cells, where 3A denotes a 4xRbHelix variant in which the 3 interface residues were substituted with alanines within all four helix sequences. n= 3.
G. Flow cytometry analysis of cell cycle phase distribution using EdU incorporation and DAPI staining of GFP-negative or GFP-positive cells binned as illustrated in Fig. S5F.
H. Comparison of G1-phase of GFP-negative or GFP-positive Rb peptide-expressing cells from Fig. S5G. n= 3. ns ($p > 0.05$), * ($p \leq 0.05$ - significant), *** ($p \leq 0.001$), and **** ($p \leq 0.0001$).
